# Bioinformatics analysis of SARS-CoV-2 RBD mutant variants and insights into antibody and ACE2 receptor binding

**DOI:** 10.1101/2021.04.03.438113

**Authors:** Prashant Ranjan, Neha, Chandra Devi, Parimal Das

## Abstract

Prevailing COVID-19 vaccines are based on the spike protein of earlier SARS-CoV-2 strain that emerged in Wuhan, China. The continuously evolving nature of SARS-CoV-2 resulting emergence of new variants raises the risk of immune absconds. During the last few months, several RBD (receptor-binding domain) variants have been reported to affect the vaccine efficacy considerably. Soon after reporting of a new double mutant variant (L452R & E484Q) in India, the country facing a deadlier second wave of infections which prompts researchers to suspects this variant to be accountable. To address the relevant concerns about this new variant affecting vaccine efficacy, we performed molecular simulation dynamics based structural analysis of spike protein of double mutant (L452R & E484Q) along with K417G variants and earlier reported RBD variants and found structural changes in RBD region after comparing with the wild type. Comparison of the binding affinity of the double mutant and earlier reported RBD variant for ACE2 (angiotensin 2 altered enzymes) receptor and CR3022 antibody with the wild-type strain revealed the lowest binding affinity of the double mutant for CR3022 among all other variants. These findings suggest that the newly emerged double mutant could significantly reduce the impact of the current vaccine which threatens the protective efficacy of current vaccine therapy.

## Introduction

COVID-19, a serious and continuously spreading pandemic affecting the world, creates severe ailments and apparently everlasting health problems. Global coronavirus virus death increases significantly. Possibilities of a wind up of this outbreak are developing adequate interventions. While Monoclonal antibody (mAb) therapy has gained emergency use approval; a few vaccines have exhibited potential & defensive effects upon COVID-19, mostly targeting the trimeric spike glycoprotein, which is involved in host cell interaction and passage to cell entry as well as the essential target for neutralizing antibodies. Essentially those were aimed against the earlier SARS-CoV-2 strain that emerged in 2019 in Wuhan China (Korber*et al*., 2020; Chen *et al*., 2020). Due to the perceived ease of transmission and expansive mutations in spike proteins, the speedy evolution of new variants of SARS-CoV-2 is of high concern. A more contagious form of the virus is spread across the country. According to recent reports by the BBC, Hindustan Times, and Irish Times, several reports of feasible re-infection have been depicted in recent months. Globally, deaths are rising again, on average around 12,000 per day, and new cases are also increasing, excelling at 700,000 per day. In Brazil, where there are about 3,000 deaths per day, or in recent weeks, a quarter of the world’s lives were lost. Situation is equally grim in India, where cases have risen in the weeks of February. The Asian nation has disclosed >14 million cases of Covid and over 174,300 deaths so far. A new variant with two mutations - E484Q and L452R, called B.1.617 initially reported in India was named “double mutant”. This new variant is believed to be creating the latest wave of infections in India, making it the second most affected country in the world, surpassing Brazil. India was returning to a deadlier new wave of infections after reporting a double mutation. This new variant was found in >200 samples from Maharashtra only, which became the first hot spot region during the second national wave that started in the last few months. By around 7 April, Maharashtra alone accounted for more than half of the new cases filed in the country, convincing experts to doubt that the variant might be blamed. Approximately, more than half of the districts in Maharashtra now have a large no. of cases than the high rate in September. Nagpur, which has returned to lockdown, records 67 new cases over 100,000 people per day, more than thrice the statewide rate. These cases are doubling every 10 days. In India, over 180,000 new infections occurred in a 24-hour period in the previous week, taking the total number of cases to over 13.9 million. For better management of this second wave, we need to re-evaluate the threshold efficacy of the vaccine explore the functional aspects or potential efficacy of vaccine therapy currently available, on the dual mutant variant and thereby look for strategic change/s for better efficacious vaccine in future. It was noted that several mutations of the receptor-binding domain (RBD), are essential for the interaction of Human angiotensin 2 altered enzymes (ACE2) (Yan *et al*. 2020) and antibodies, as well as region that neutralizes antibodies. It is therefore imperative to understand up to what extent mutations interrelated with SARS-COV-2 affect the vaccination. Taking these into account both vaccination as well natural infections, several reports deal with the outcome of these variants on antibody ligation and function. The RBD can exist in two conformities, alluded to as “up” and “down” i.e. receptor accessible and receptor in-accessible (Wrapp *et al*. 2020). The *in silico* investigation revealed that ACE2 and potential antibodies binds in a similar area on the spike protein (Hwang *et al*. 2006, Sui *et al*. 2004). An antibody becomes very effective when it forestalling viral spread by impeding the ACE2 binding site in the RBD. CR3022 antibody showed the most elevated binding affinity with SARS-CoV-2 S-protein RBD (Hussain *et al*. 2020, Huo*et al*. 2020).

Based on the up to date literature survey, we retrieved 28 different spike protein variants and out of these 28 variants, 12 variants belong to RBD region only. Here, we report RBD variants that affect the ability of CR3022 and ACE2 to bind with the SARS-CoV-2 RBD through *insilico* analysis.

## Material and methods

### Retrieval of crystal structures

Crystalstructuresof spike protein (PDBID-7AD1), ACE2 (PDBID-6ACG) and antibody CR3022 (PDBID 6YLA) were retrieved from PDB RCSB (https://www.rcsb.org/). All water molecules and hetro-atoms were removed by using Discovery studio visualization software (BIOVIA 2020). (http://accelrys.com/products/collaborative-science/biovia-discovery-studio/visualization-download.php).

### Homology modeling and Energy minimization

Based on high similarity, 7AD1 (crystal structure of SARS-CoV-2) was selected as template for homology modeling of RBD mutant variants using the SWISS-MODEL (Lyskov and Gray 2008).Energy minimization and structural analysis of RBD mutant variants were done with UCSF Chimera (Pettersen *et al*. 2004). Evaluation of the modeled structure was done by PDB-Sum (http://www.ebi.ac.uk/thornton-srv/databases/cgi-bin/pdbsum/GetPage.pl?pdbcode=index.html).

### Docking analysis

Docking of RBD mutant variants with selected targets (ACE2 receptor and antibody structure CR3022) was carried out by PatchDock server (Ranjan *et al*. 2020) by choosing parameter RMSD esteem 4.0 and complex type as default. Docking investigation was based on geometric shape complementarity score. Higher score indicates higher binding affinity. Outcome of the results is based on the docking scores and interaction at the RBD regions. Protein-protein and antibody-protein interactions were visualized by LigPlot plus v2.2 (Wallace *et al*., 1995).

Molecular interactions of antibody CR3022 and ACE2 receptor with RBD variants were performed byantibody script under antibody loop numbering scheme i.e. KABAT Scheme and DIMPLOT script algorithm package built into LigPlot plus v2.2 respectively.

### Molecular Dynamics Simulation

The equilibrium and the dynamic behavior of wild and mutant variants of RBD Spike protein was studied by using GROMACS (Hess, et al., 2008).

MD simulation brings about time-dependent conformational changes and adjustment of protein, which opens to the alteration in unique nature after establishment of mutation in protein. We used GROMOS96 54a7 force field (Pandey, et al., 2020) for MD simulation study. We added solvent water around protein to facilitate from spc216.gro as a non exclusive equilibrated 3-point dissolvable water model in a dodecahedron. Here, we kept the protein in the centre in at least 1.0 nm from the case edges.

Further, steepest descent algorithm was utilized for energy minimization, to remove the steric conflicts and unstable conformations. Further we equilibrate the system via NVT ensemble (constant Number of particles, Volume and Temperature) and NPT ensemble (constant Number of particles, Pressure and Temperature). After achieving equilibrium process, we moved for MD run to 10ns. Data analysis was done by Gromacs tools i.e. gmx rms for RMSD (Root Mean Square Deviation), gmx rmsf for RMSF (Root Mean Square Fluctuation), gmx gyrate for radius of gyration (Rg), gmx hbond for H-bond (for intra-protein H-bonds and for H-bonds between protein and water), and gmx sasa for SASA (solvent accessible surface). We further used GRACE software for data visualization.

## Results

We have retrieved 28 variant mutants **(Figure1)** in spike protein from literature survey. We found 12 variants/mutants in RBD region. RBD region is important for ACE2 and Antibody interactions. We have done structural analysis of all 12 mutant variants and compared with wild type and found seven mutant variants (F486L, Q493N, double mutant (L452R & E484Q), R408I, L455Y, K417G and E484K) have structural changes in RBD region **(Figure2)**. We analyzed interactions between RBD variants and ACE2 receptor. Moreover, we checked the interactions between antibody and RBD variants too. It was found that seven structurally changed variants (F486L, Q493N, Indian double mutant (L452R & E484Q), R408I, L455Y, K417G and E484K) have high docking score against ACE2 receptor compared with wild type and less docking score against antibody (CR3022) unlike wild type **(Table2)**. High docking score signifies high binding affinity and low docking score signifies low binding affinity. Out of seven variants, double mutants (Double mutant< Q493N<E484K<K486L<L455Y<R408I<K417G) have lowest binding energy against antibody. Molecular interactions of antibody and ACE2 receptor with RBD variants are depicted in **supplementary file1**. A few RBD variants already shown to affect the vaccine efficacy as documented earlier by wet lab and dry lab results **(Table1)**, however, the vaccine efficacy against the double mutant and K417G variants is yet to be elucidated. Our *in silico* study suggests that the double mutant and K417G variants may affect the vaccine efficacy.

**Table1.**
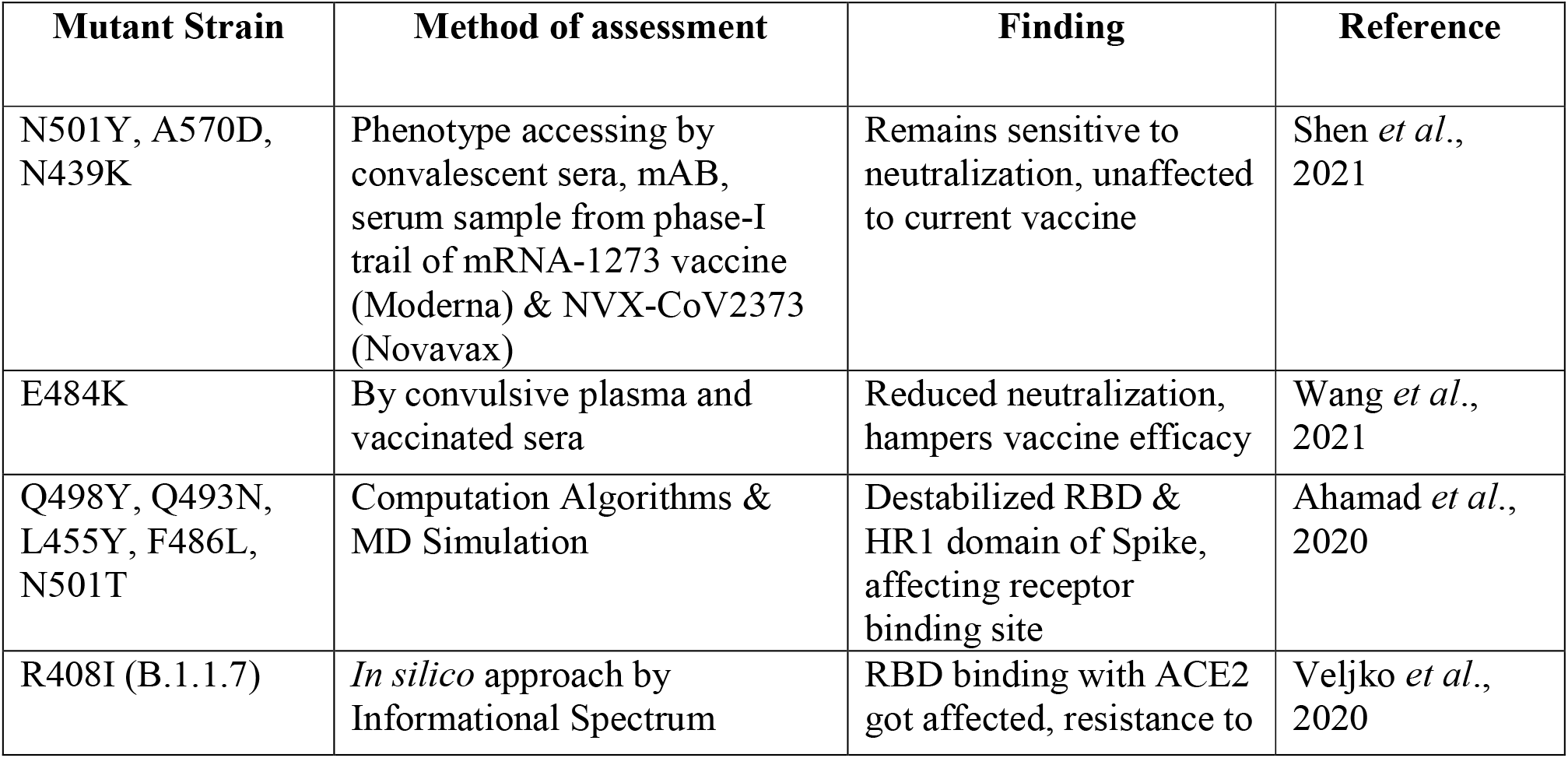
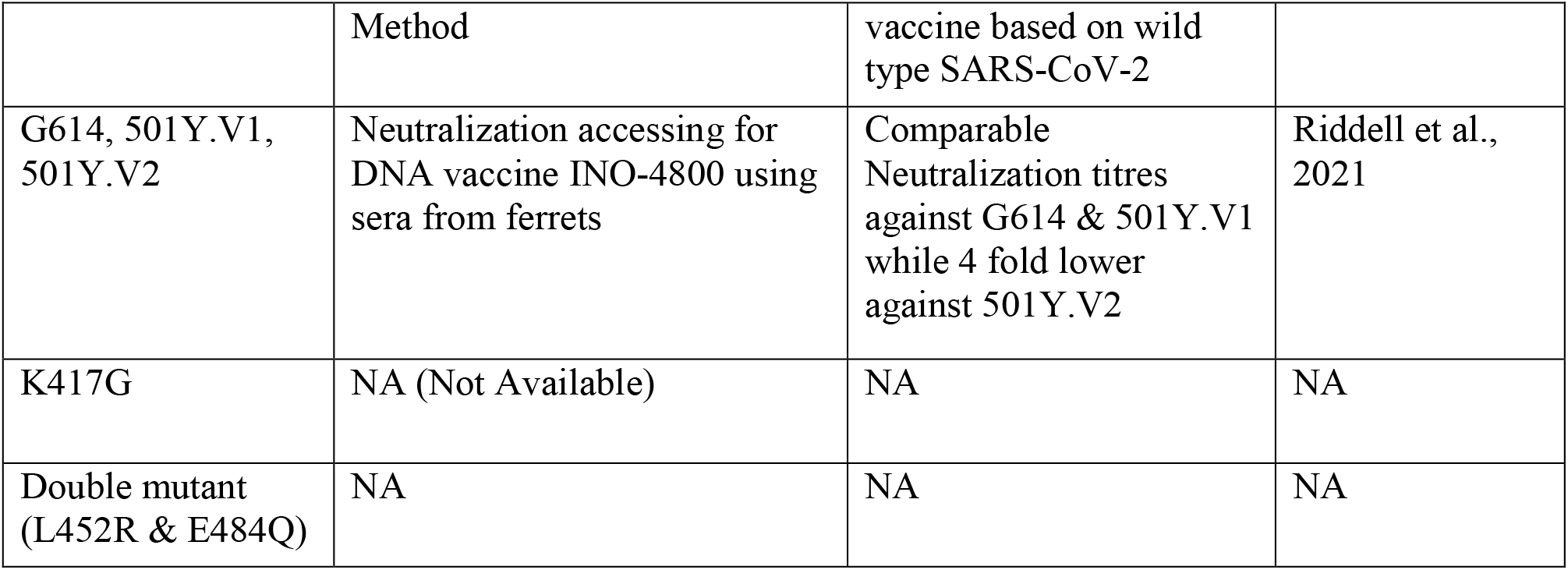
Table showing studies related to vaccine efficacy hampered by RBD based variants detected by different methodology.

**Table2.**
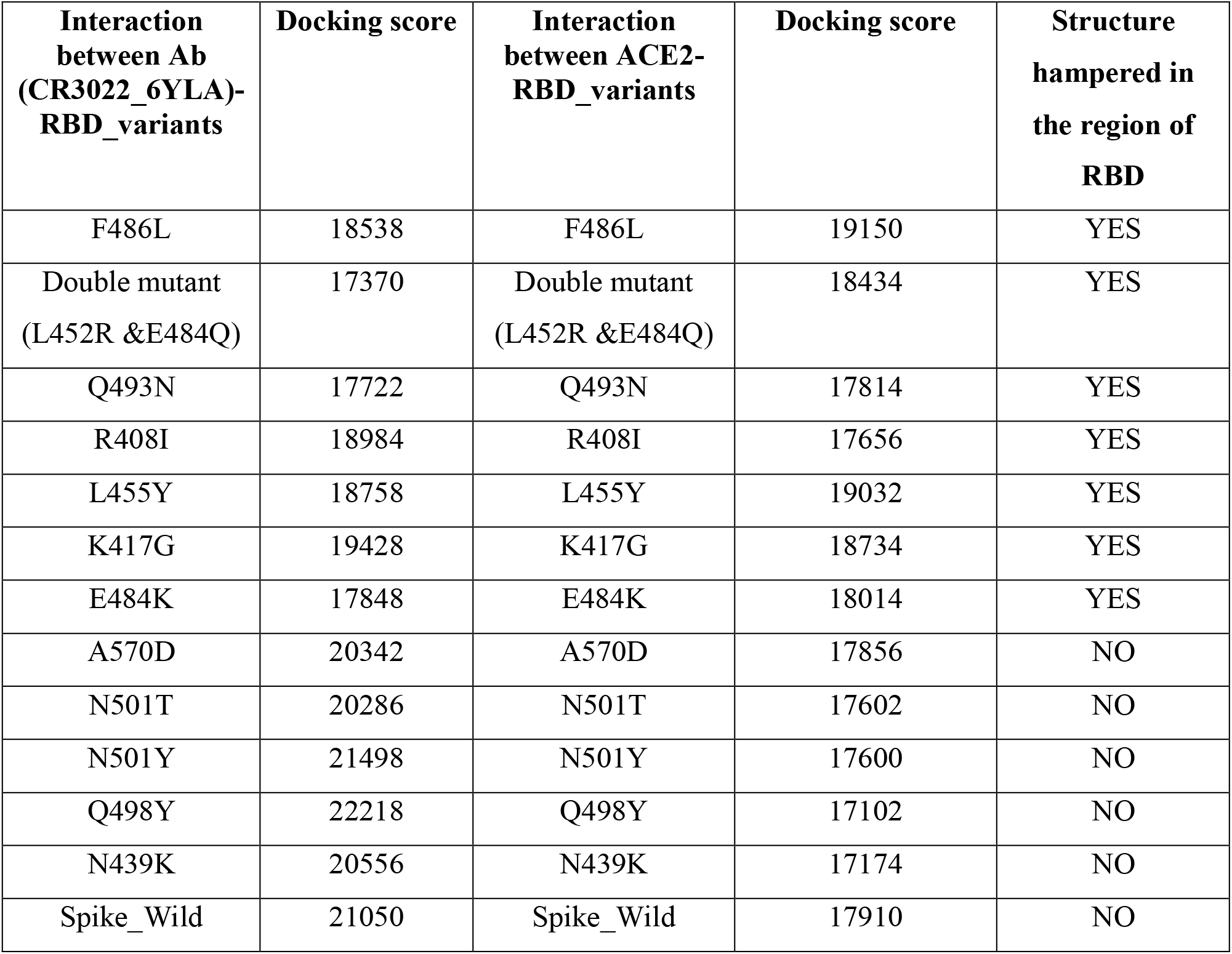
Prediction of RBD based variants interaction with antibody and ACE2 receptor.

| **Interaction between Ab (CR3022_6YLA)-RBD_variants** | **Docking score** | **Interaction between ACE2-RBD_variants** | **Docking score** | **Structure hampered in the region of RBD** |
| --- | --- | --- | --- | --- |
| F486L | 18538 | F486L | 19150 | YES |
| Double mutant (L452R &E484Q) | 17370 | Double mutant (L452R &E484Q) | 18434 | YES |
| Q493N | 17722 | Q493N | 17814 | YES |
| R408I | 18984 | R408I | 17656 | YES |
| L455Y | 18758 | L455Y | 19032 | YES |
| K417G | 19428 | K417G | 18734 | YES |
| E484K | 17848 | E484K | 18014 | YES |
| A570D | 20342 | A570D | 17856 | NO |
| N501T | 20286 | N501T | 17602 | NO |
| N501Y | 21498 | N501Y | 17600 | NO |
| Q498Y | 22218 | Q498Y | 17102 | NO |
| N439K | 20556 | N439K | 17174 | NO |
| Spike_Wild | 21050 | Spike_Wild | 17910 | NO |

**Figure1.**
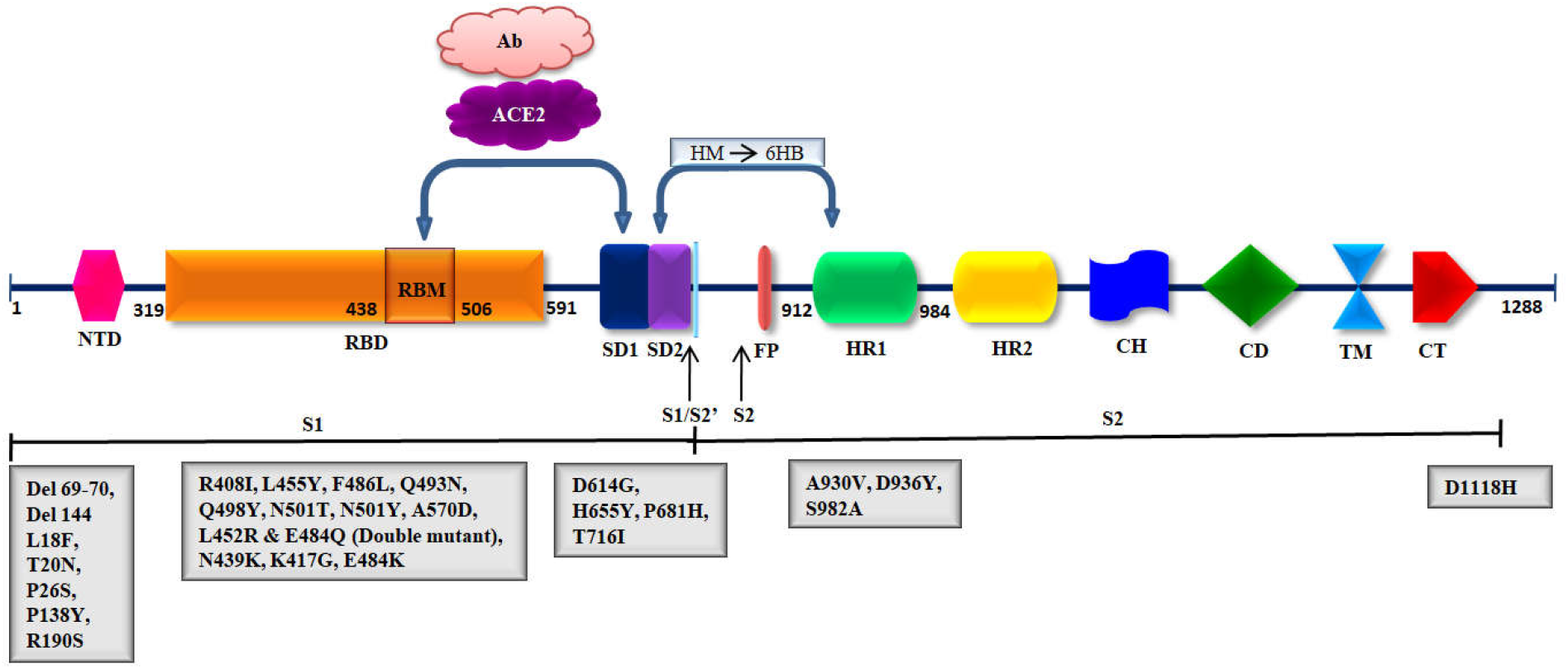
Schematic representation of SARS-CoV-2 spike glycoprotein along with depiction of ACE2 and Antibody binding on RBD. NTD: N-terminal domain, RBD: receptor-binding domain, RBM: receptor-binding motif, SD1 and SD1: Subdomain1 and 2, S1 and S2: Protease Cleavage Site 1 and 2, FP: fusion peptide, HR1 and HR2 heptad-repeat regions 1 and 2, CH: Central Helix, CD: Connector Domain, TM transmembrane region, CT cytoplasmic tail, HM: Homotrimeric assembly, 6HB: Six Helix bundle.

**Figure2.**
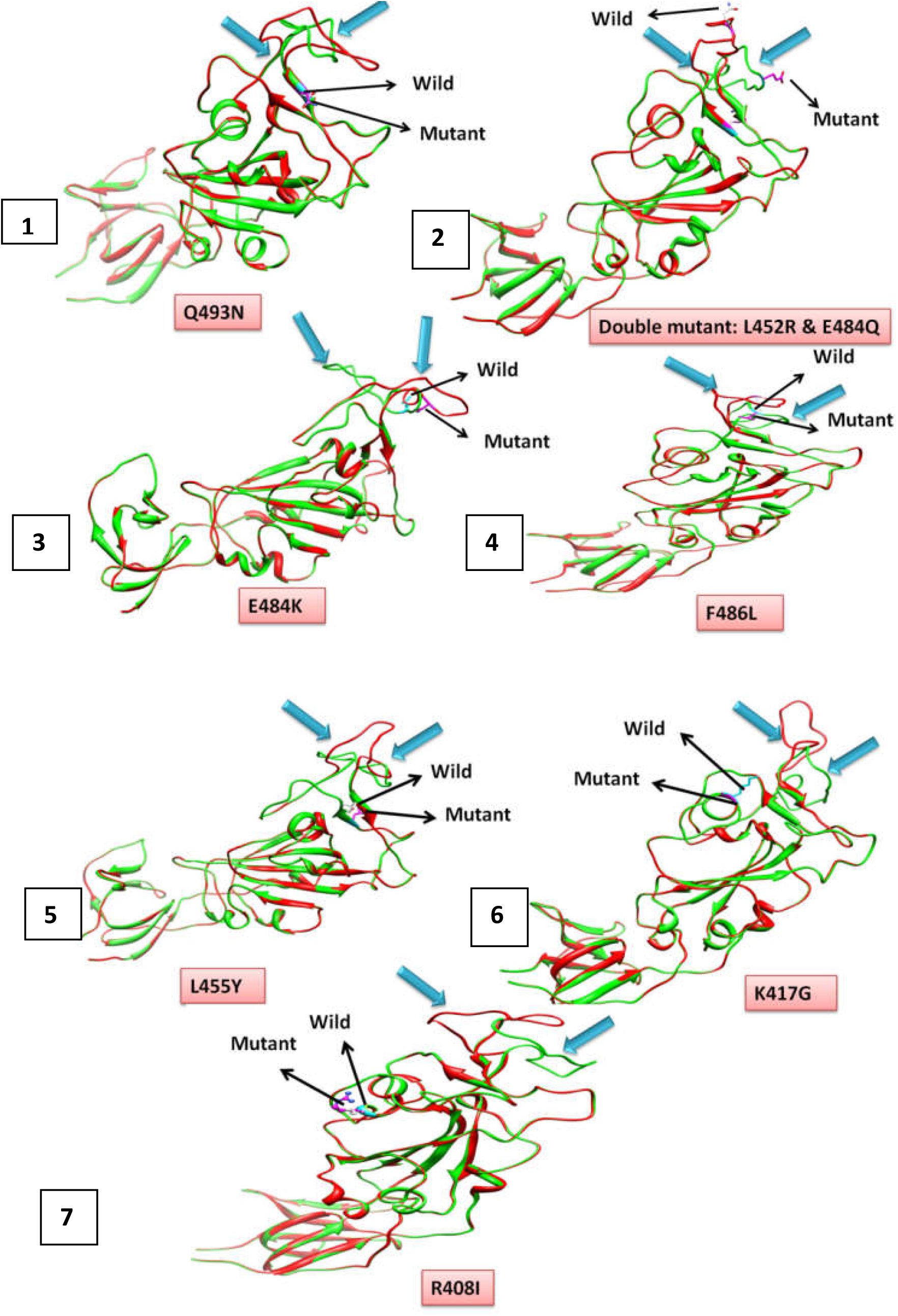
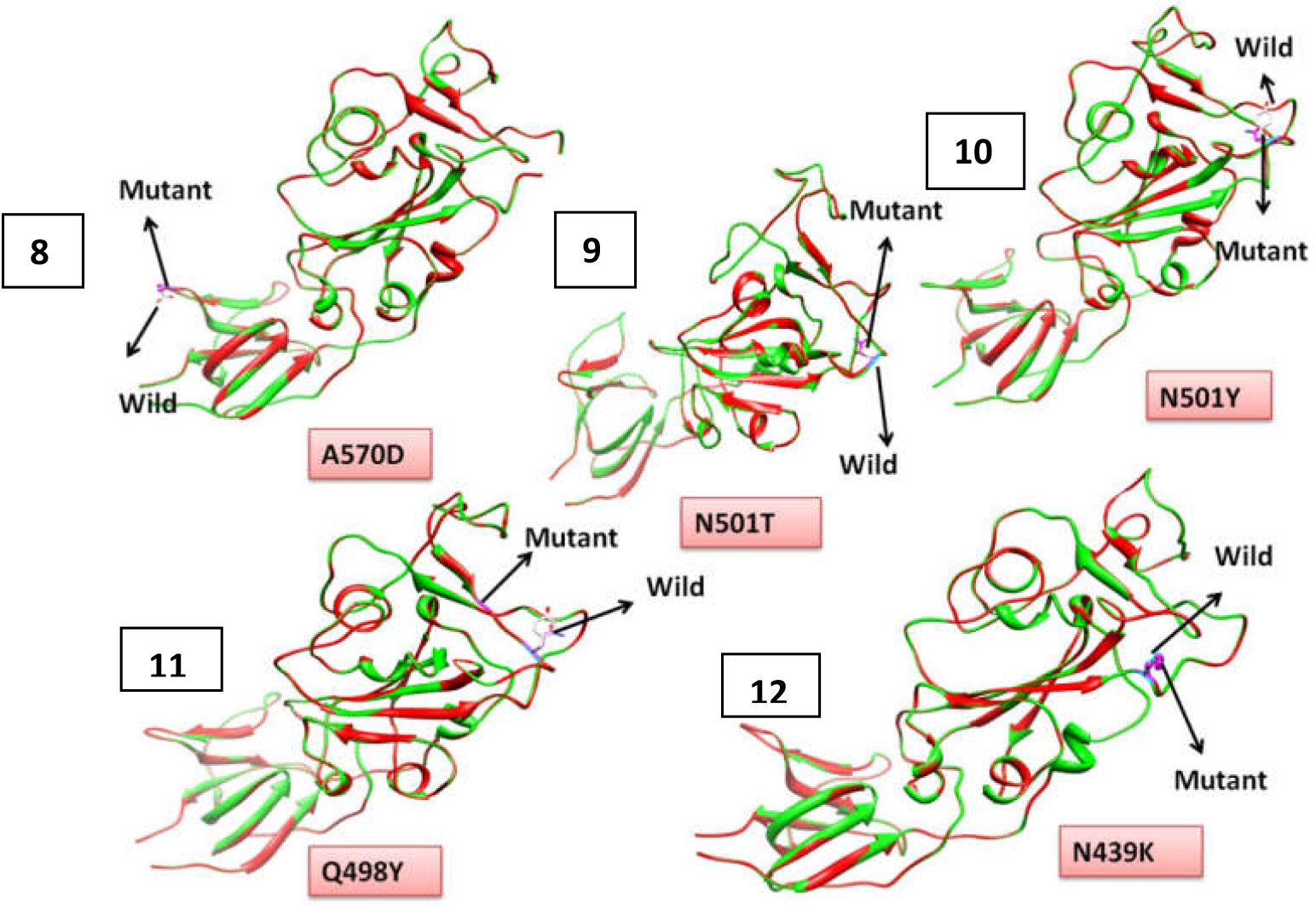
Structural superposition of RBD based variants with wild type: 1-7 represents seven RBD mutant variants depicting structural changes when compared with wild type. 8-12 shows five RBD mutant variants do not have changes on RBD region. Green color indicates wild type and red color indicates mutant.

### Molecular Dynamics Simulations

To examine the dynamic behavior, MD simulation runs for 10 ns to contemplate the structural stability of RBD mutant variants (F486L, Q493N, Indian double mutant (L452R & E484Q), R408I, L455Y, K417G and E484K) as compared to wild type. Various parameters studied all through the simulation trajectory, such as RMSD, Rg, RMSF, SASA, total number of intra-molecular hydrogen bonds of protein and H-bond between protein and water with the time dependent function of MD to examine the functional and structural impact of a mutant on wild protein.

The RMSD and RMSF **(Figure 3&4)** of C-alpha chain atoms of all RBD mutant variants showed significant fluctuations in stability as well as in flexibility in comparison to Wild one. Rg and SASA analysis **(Figure 5&6)** depicted much fluctuation between mutant and wild type. The fluctuations of hydrogen bonds are easily shown in all RBD mutant variants as compared to wild type (total intra-molecular H-bond and H-bond between protein and water) **(Table3)**. These molecular simulation data showed hampered structural stability and complexity of all seven RBD mutant variants as compared to wild type.

**Table3.** Comparative studies of total intra-molecular H-bond and H-bond between protein and waterof all RBD mutant variants along with wild type. *HD: Hydrogen Donor HA: Hydrogen Acceptor.

| Variants | Total-Intramolecular hydrogen bonding |  | H-bond between protein and water |  |
| --- | --- | --- | --- | --- |
|  | HD* | HA* | HD* | HA* |
| **Wild Type** | 397 | 767 | 20501 | 20871 |
| **Double mutant** | 401 | 770 | 23601 | 23970 |
| **E484K** | 398 | 766 | 26606 | 26974 |
| **F486L** | 397 | 767 | 23200 | 23570 |
| **K417G** | 396 | 766 | 28484 | 28854 |
| **L455Y** | 398 | 768 | 25553 | 25923 |
| **Q493N** | 397 | 767 | 25576 | 25946 |
| **R408I** | 394 | 764 | 25532 | 25902 |

**Figure3.**
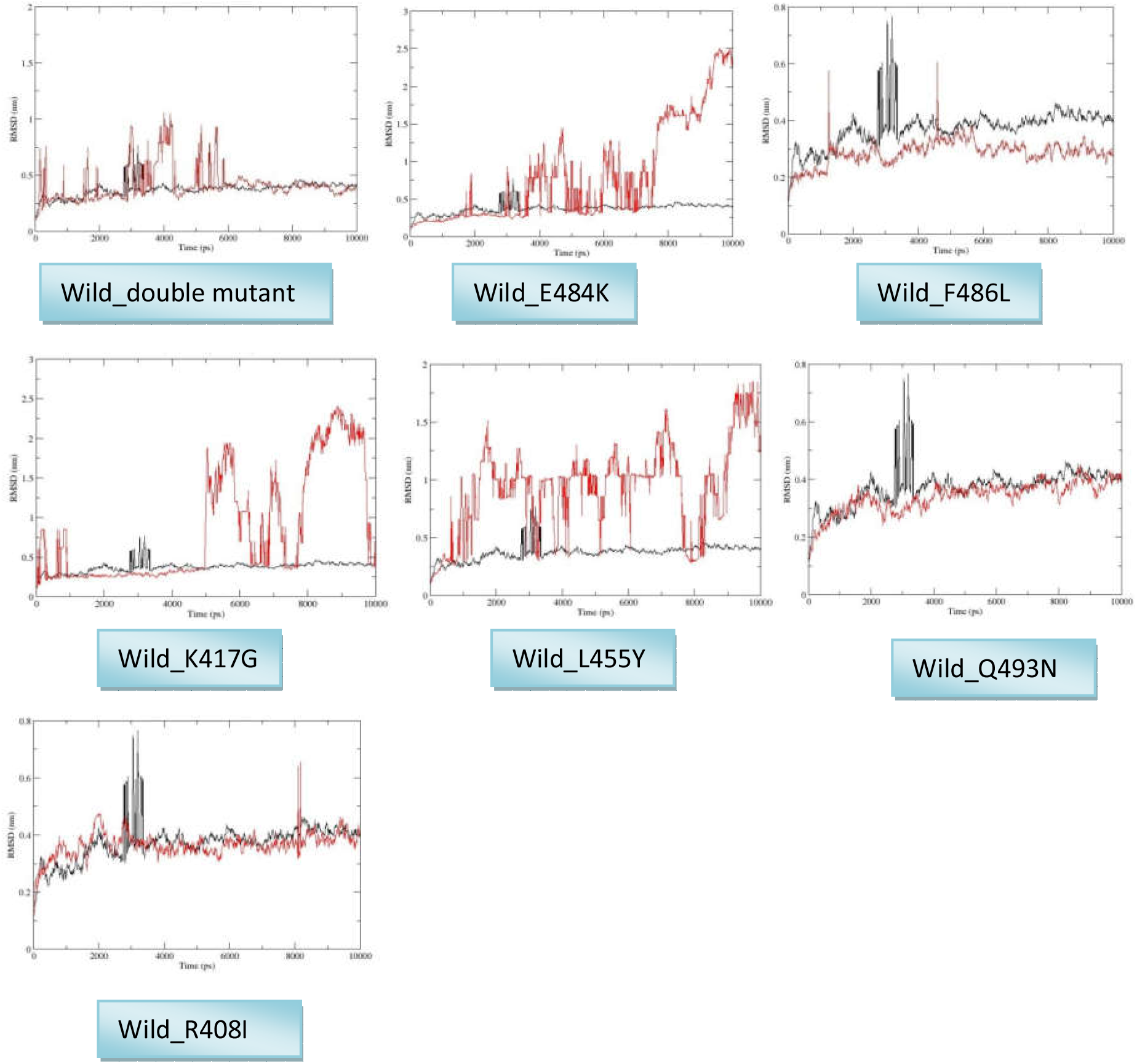
RMSD analysis of RBD mutant variants along their wild type over 10 ns simulation. Black dot plot showed wild type where as red indicates mutant.

**Figure4.**
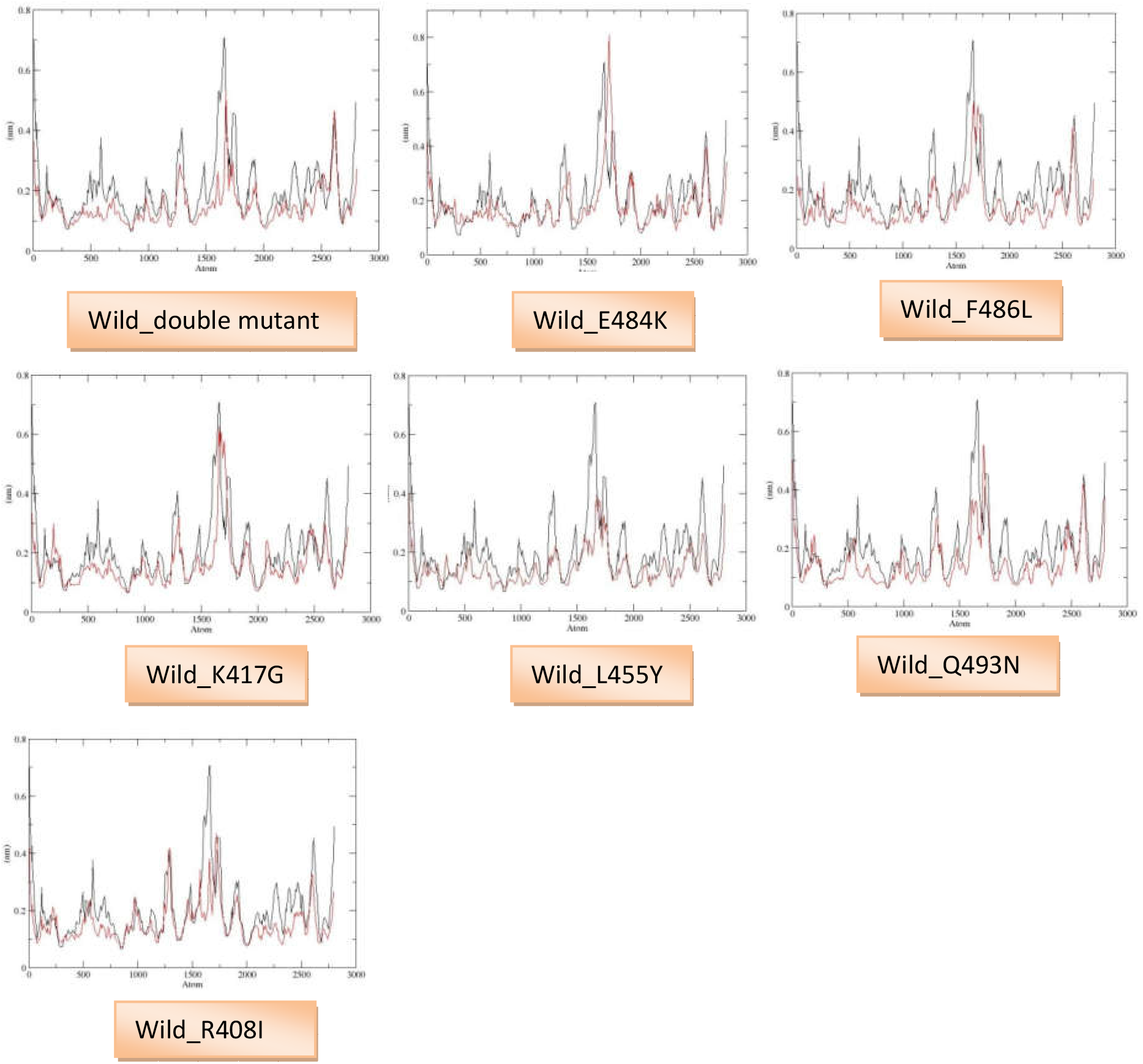
RMSF analysis of RBD mutant variants along their wild type over 10 ns simulation. Black dot plot showed wild type where as red indicates mutant.

**Figure5.**
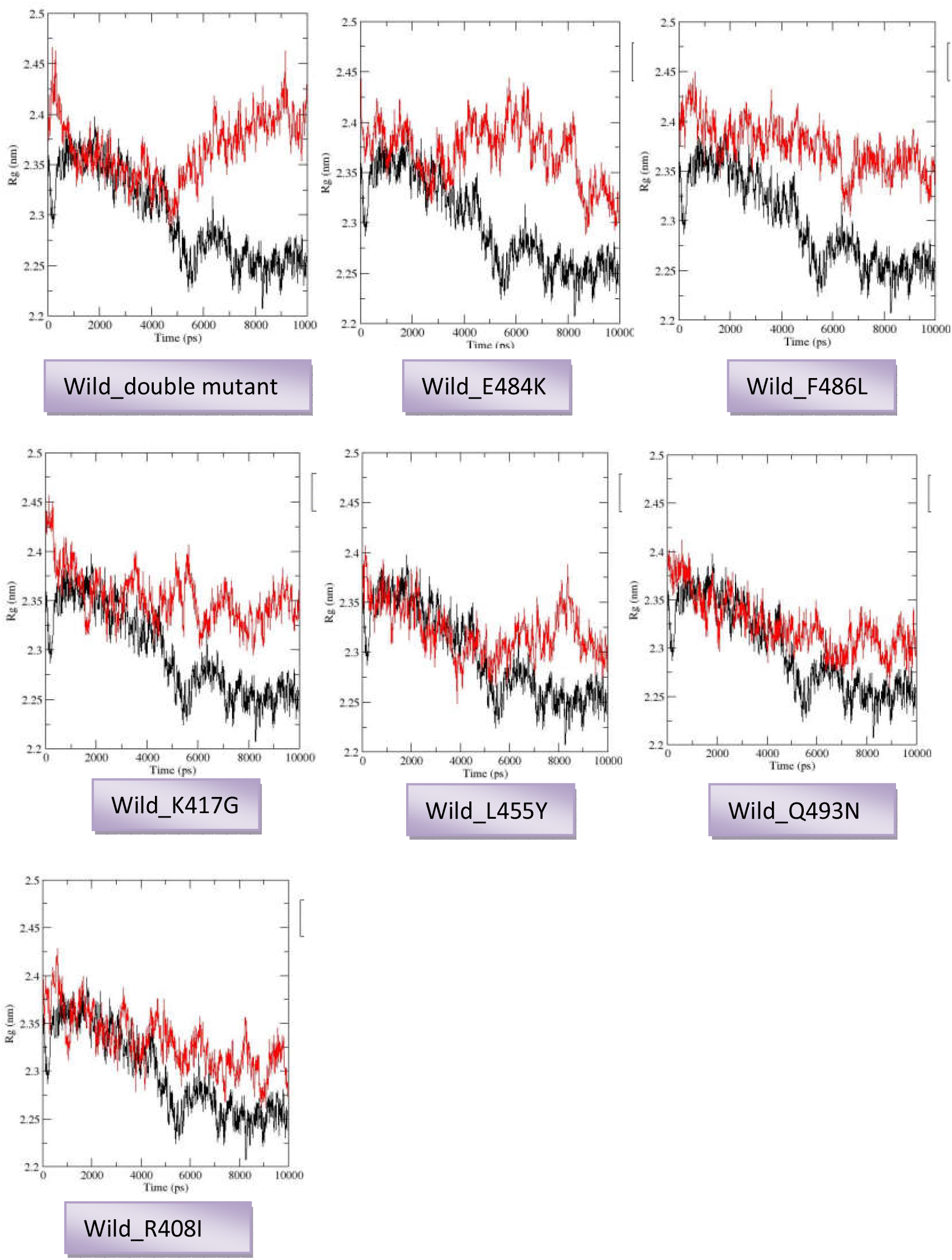
Rg analysis of RBD mutant variants along their wild type over 10 ns simulation. Black dot plot showed wild type where as red indicates mutant.

**Figure6.**
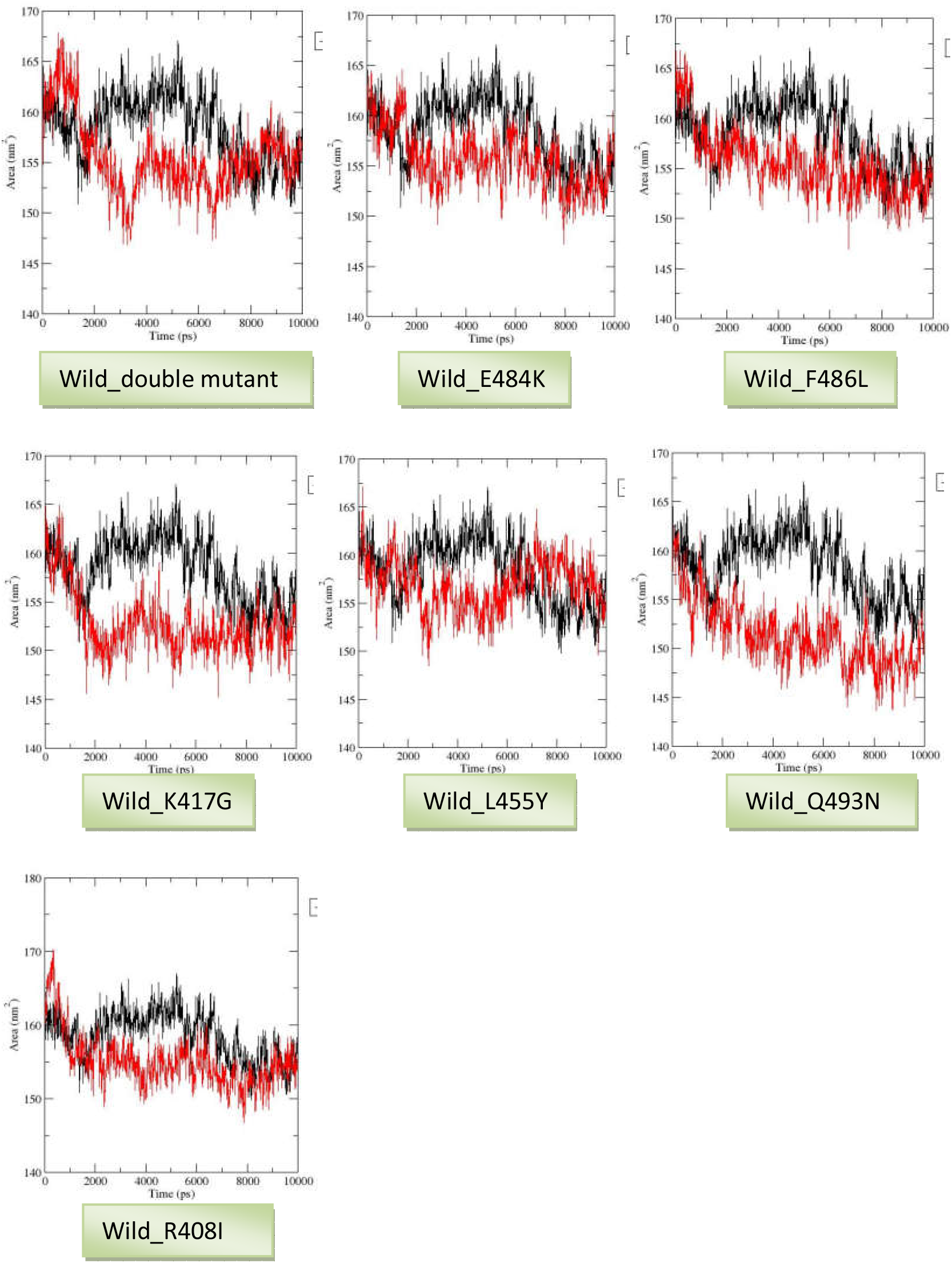
SASA analysis of RBD mutant variants along their wild type over 10 ns simulation.Black dot plot showed wild type where as red indicates mutant.

## Discussion

Continuing emerging variant of SARS-Cov-2 resulting compromised vaccine induced immunity. The 20I / 501Y.V1 variant of the lineage B.1.1.7, first discovered in the UK, has eight major mutations in the spike genes that may affect vaccine efficiency, antibody therapy, and the threat of re-infection. In addition to remaining susceptible to antibody neutralization, the B.1.1.7 variant does not seem to be a major burden for available vaccines (Shen *et al*., 2021; Muik*et al*., 2021).

B.1.351, a variant first encountered in South Africa, is of greater concern that this variant is incompliant to NTD mAbs neutralization, mainly due to E484K mutations. In addition, B.1.351 was more opposing to neutralization by convulsive plasma (9.4-fold) and vaccinated sera (10.3– 12.4-fold) (Wang *et al*., 2021). The SARS-CoV-2 P.1, the Brazilian variant of B.1.1.28 lineage, has 10 mutations in spike gene viz. D614G, T20N, D138Y, L18F, R190S, and P26S in the NTD and K417T, E484K and N501Y in the RBD region and H655Y within furin cleavage site. It shares mutations similar to B.1.35. P.1 on the same 3 RBD residues which are resistant to neutralization by the RBD targeted mAbs. Shared E484K mutation is the main culprit, which emerged in more than 50 lines independently along with B.1.526, recently identified in New York. A significant loss of neutralizing activity has been shown by vaccinated serum and convalescent plasma towards P.1, but the decrease is not as goodas compared to what was found against B.1.351., Accordingly, the risk of re-infection by P.1 or dropped efficacy of vaccine protection may not be severe like B.1.351 (Wang *et al*., 2021).

The mRNA-1273 vaccine’s neutralizing activity towards number of variants like B.1.351, B.1.1.7 + E484K, B.1.1.7, P.1, B.1.427 / B.1.429, D614G, 20A.EU2, 20E [EU1], N439K-D614G, and previously identified mutant in Denmark mink cluster 5 were identified and found to have the same neutrality level as Wuhan-Hu-1 (D1414) (Wu *et al*., 2021). Limited loss in antibody neutralizing activity against B.1.1.7 while significant loss against B.1.35 was shown by the AstraZeneca ChAdOx1 vaccine, thus maintaining its efficacy towards B.1.1.7 and demonstrating a major loss of efficacy against the benign version of B.1.151. Although the efficacy against B.1.1.7 was found to have retained by the BNT162b2 Pfizer / BioNTech COVID-19 vaccine. The Novavax vaccine (NVX-CoV2373) reported differential protective immunity in the clinical trials i.e 96%, 60%, and 86% against parental strain, B.1.351 and B.1.1.7, respectively (Tarke*et al*., 2021).”Double mutant” coronavirus variation with a combination of changes not seen in any other places in the world than in India according to Times of India and BBC news. Many SARS-CoV-2 variants have been detected in the last few weeks of March in India. India is experiencing a sharp rise in coronavirus infections from between March and April 2021. Based on worldometer statistics data https://www.worldometers.info/, we observed sudden enhancement of total, active, daily new cases, and death rate in the months of March and April indicates the beginning of second wave in India **(Figure 9)**. We do also observed the confirmed and active cases in all State/Union territory (UT) of India with sudden rise from March to April 2021 according to covid19 India tracker (http://www.covid19india.org) **(Figure10&11)**. Based on these studies, it can be hypothesized that the double mutant variant may also one of the causes of the sudden increment of cases in all state and UT of India. It might be possible that double mutant variant plays a major role in speedy spreading of infections in India, which we tried to explore through *in sillico* analysis.

**Figure7.**
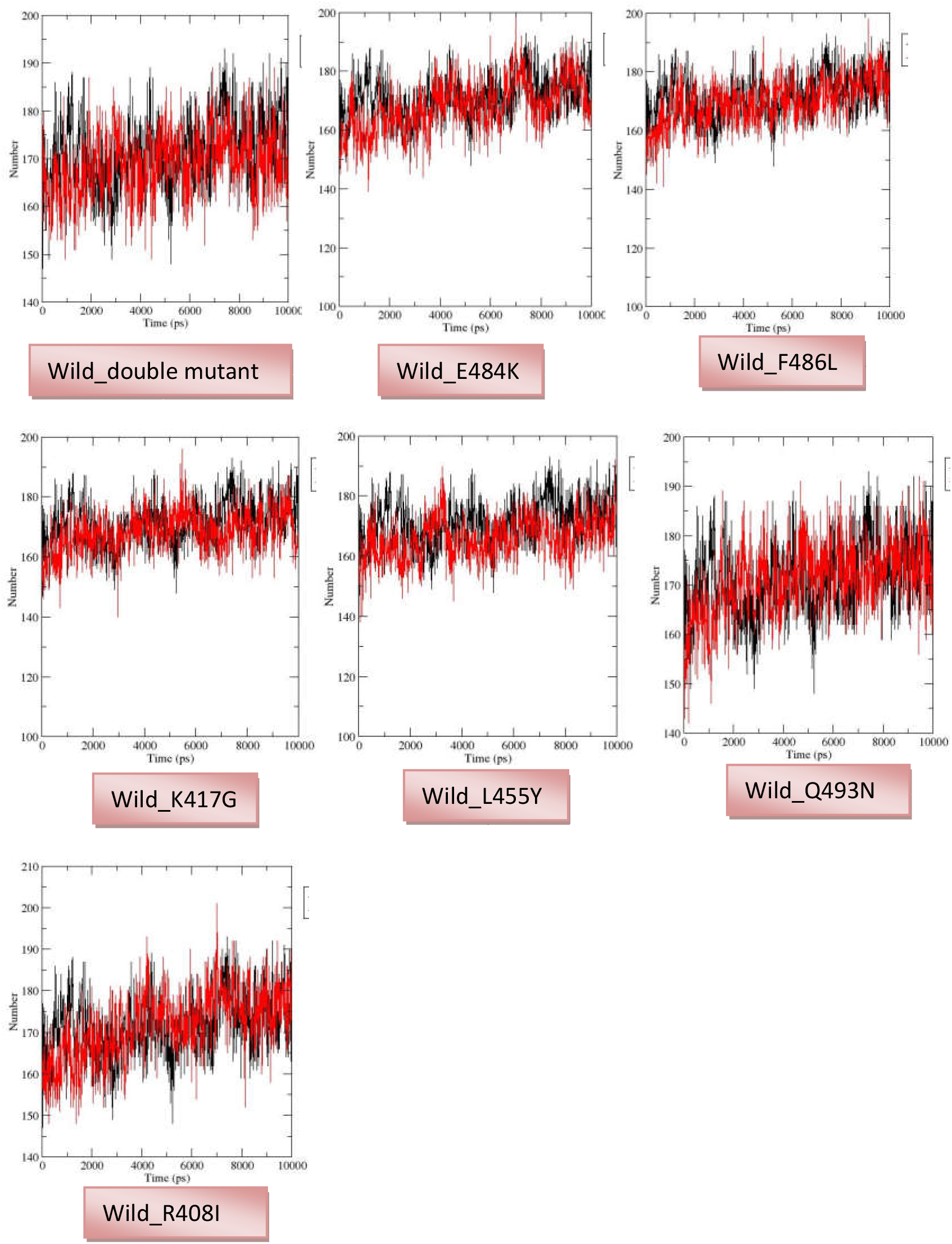
Total intra-molecular H-bonding analysis of RBD mutant variants along their wild type over 10 ns simulation. Black dot plot showed wild type where as red indicates mutant.

**Figure8.**
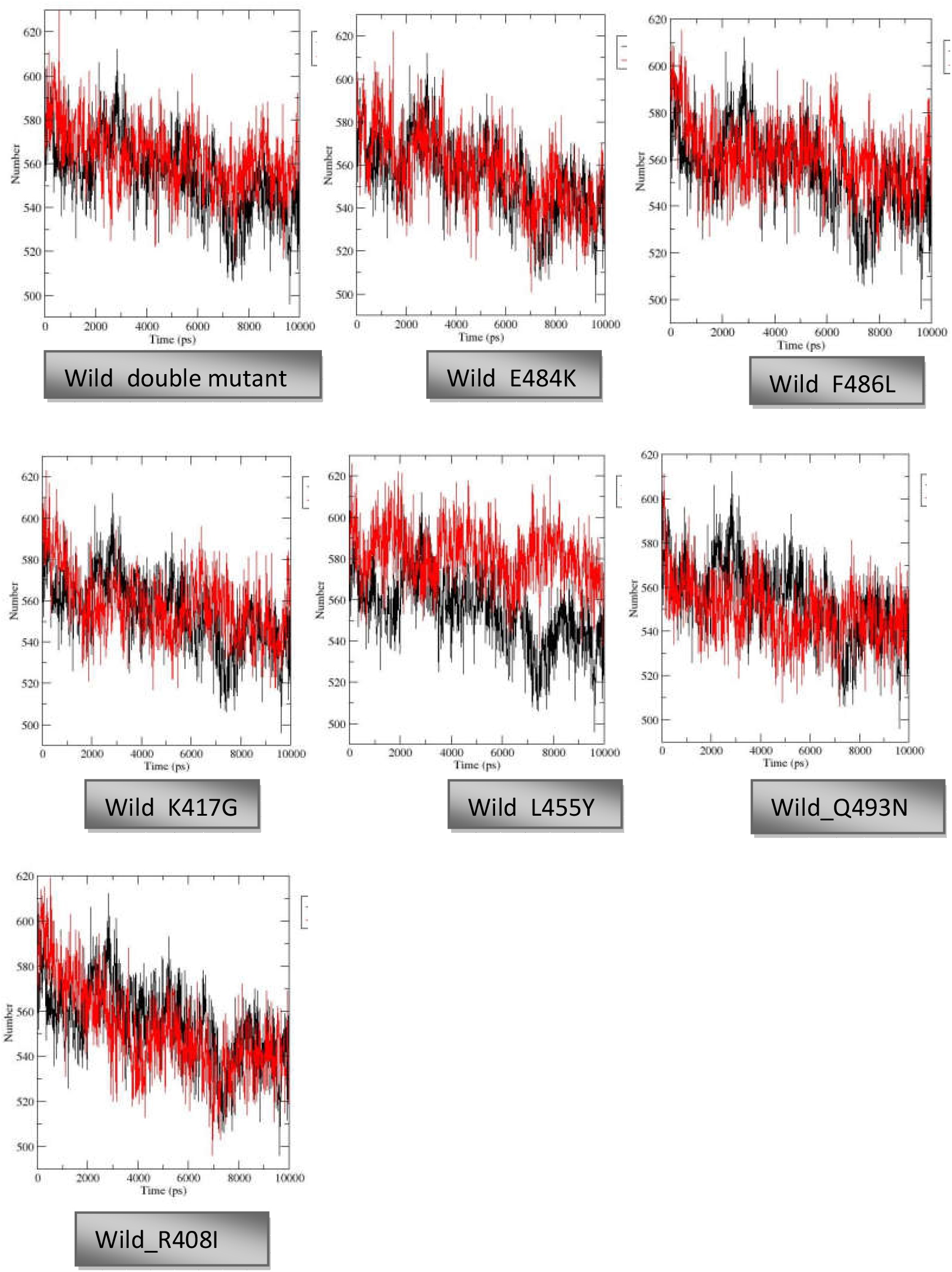
Analysis of H-bond between protein and water of RBD mutant variants along their wild type over 10 ns simulation. Black dot plot showed wild type where as red indicates mutant.

**Figure9.**
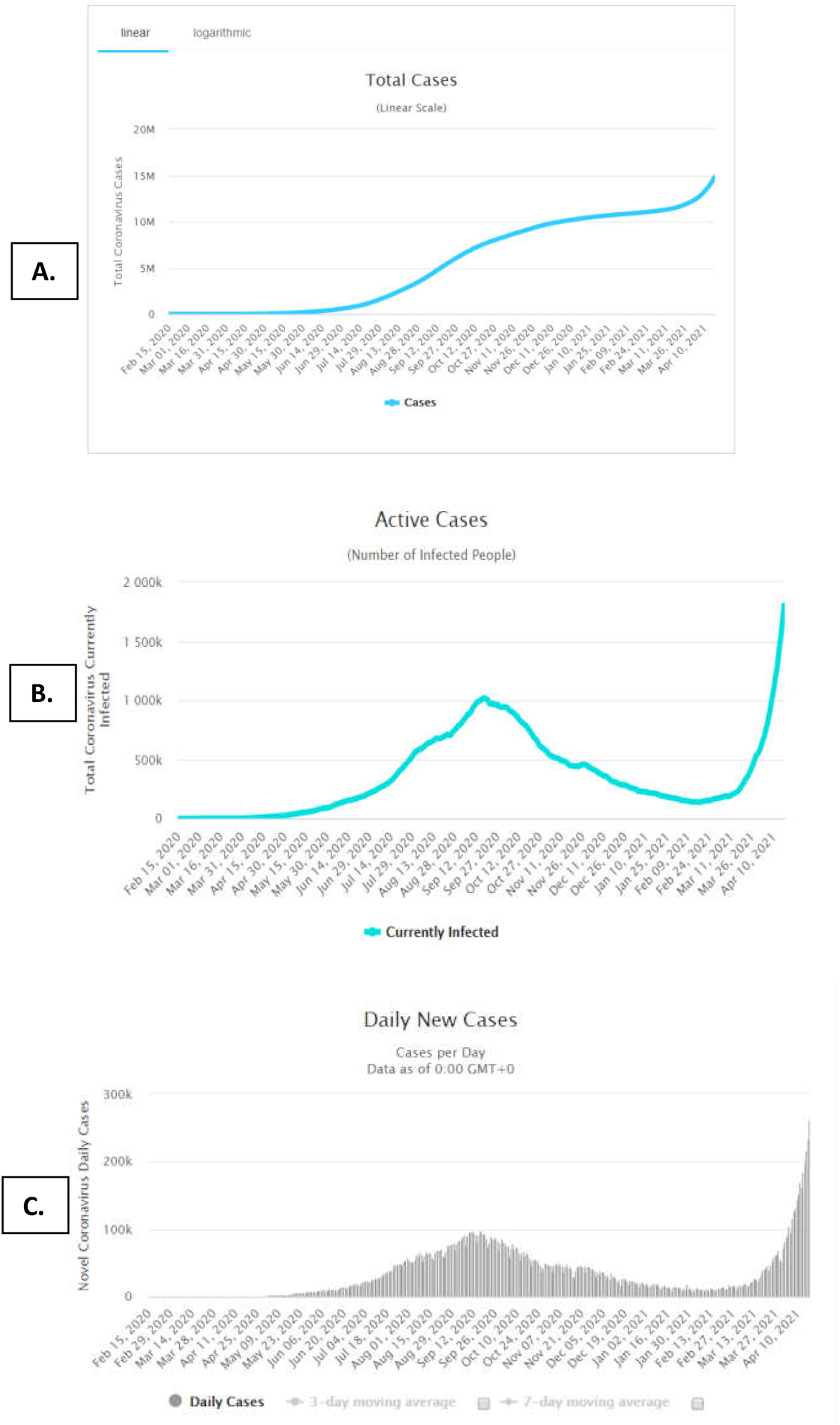
Representation of cases sudden rises in the months between March and AprilA. Total confirmed, B. Active and C. daily new cases in India. (https://www.worldometers.info/)

**Figure10.**
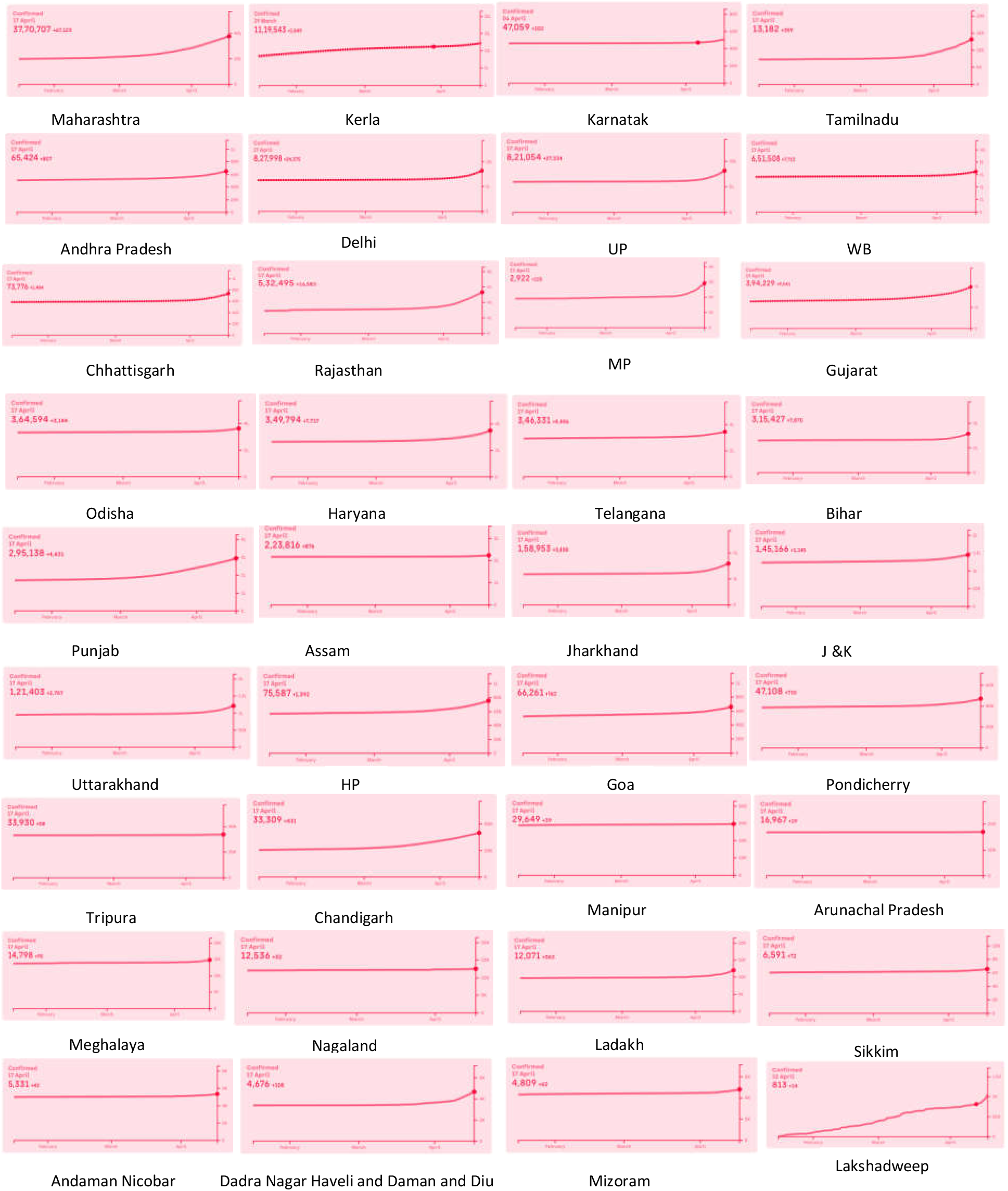
Confirmed cases reported in State/UT of India from Jan to April 2021. (http://www.covid19india.org)

**Figure11.**
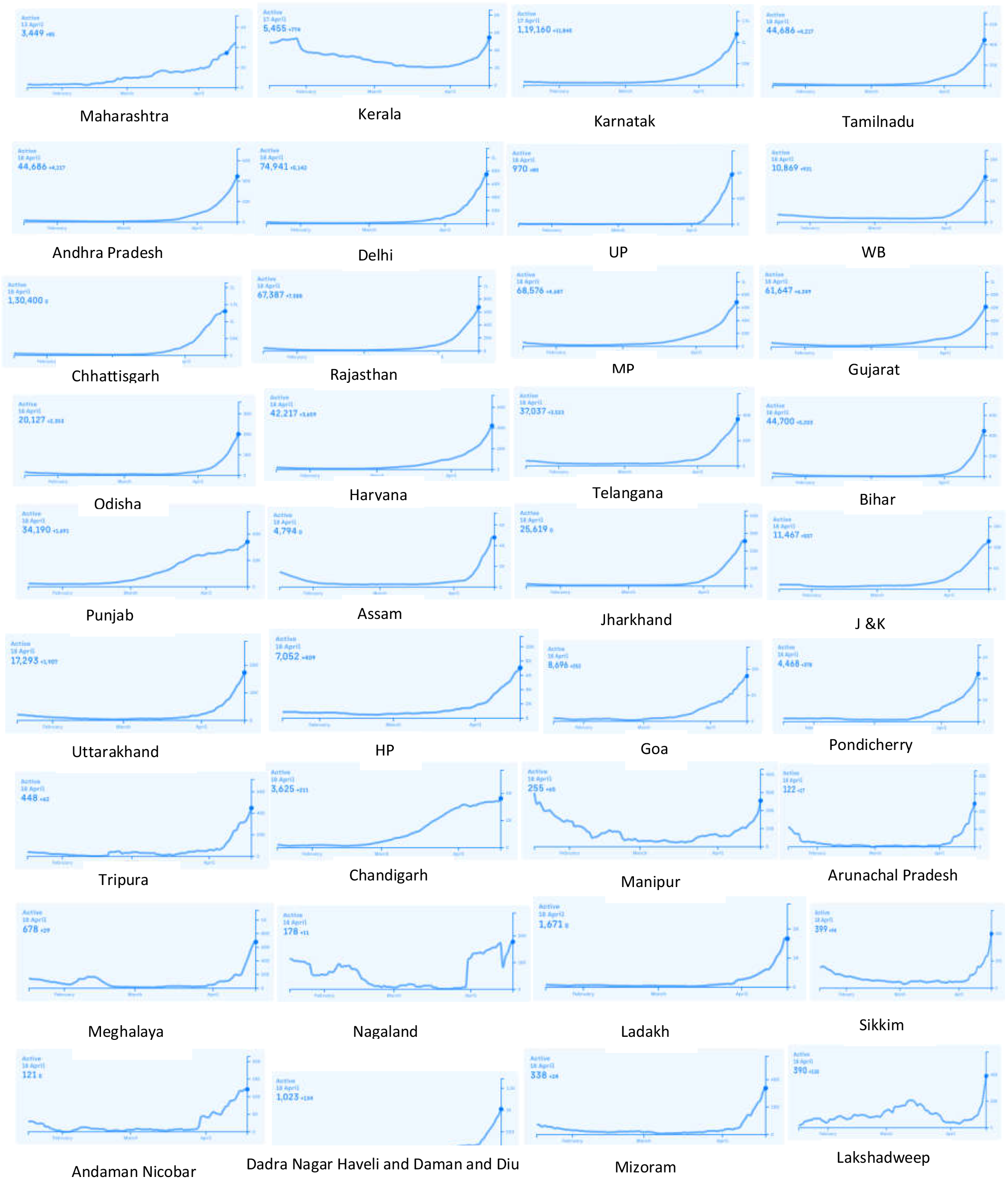
Active cases reported in State/UT of India from Jan to April 2021. (http://www.covid19india.org)

The RMSD, RMSF, Rg, SASA, Intramolecular H-bond and H-bond between protein and water molecules were plotted to analyze the stability as well as flexibility of structural hampered mutant RBD variants. Comparison of wild protein with mutant protein, higher RMSD fluctuations observed in all variants except Q493N, K408I. In case of Q493N and K408I less fluctuation was noted. The RMSD output showed that the protein stability can be influenced. We observed that the RMSF values are lower as compared to wild and it confirms the compressed behavior of mutant trajectory. We noticed higher value of Rg in case of all mutants which indicates that the compactness of protein can be lower. High fluctuations of SASA results revealed that the protein structure and consequently protein function might be hampered. The fluctuations of total intra-molecular H-bond and H-bond between protein and water showed in all structural hampered RBD mutant variants which indicates the rigidity of protein may be influenced (Hubbard *et al*., 2010).

Previous study has disclosed that the residues F486, L455, Q493, and N501 in the RBD spike protein form a major binding domain for the human ACE2 receptor (Wan *et al*., 2020). A few mutants’ viz.L455Y, Q493N, R408I, Q498Y, F486L, N501T within the RBD region (319-591), and D936Y & A930V within HR1 site (912-984) have also been studied by *in silico* analysis to investigate the basic structure of spike glycoprotein. After comparing MD simulations in mutants and WT, a significant destabilizing outcome of mutations on the HR1 and RBD domains was revealed. Researchers revealed compromised stability of the overall spike protein structures by investigating the effect of framed mutations, before binding to the receptor (Ahamad *et al*., 2020).

## Conclusion

In this severe COVID-19 pandemic, in silico studies can be used to stimulate the proceeding of development of therapeutic agents for the treatment of disease. In this study, we performed molecular docking and simulation based screening of recently emerged double mutant variant and previously reported RBD variant of COVID-19 to compare the binding and functional stability of spike proteins. Results of the present study suggest that the new Indian strain double mutant and K417G within the receptor-binding site could reduce the vaccine efficacy by affecting the SARS-CoV-2 interaction with the CR3022 antibody and ACE2 receptor. We have examined the impact of double mutants and earlier reported RBD variants on the spike glycoprotein’s structural stability by *in silico* analysis along with molecular simulation data and found structural alteration in the RBD domain in seven mutant variants. Further molecular interaction study of CR3022 antibody and ACE2 receptor with the RBD variants and comparison with wild type strain revealed the reduced binding affinity of double mutant with antibody, besides double mutant found to have the lowest affinity among all the RBD variants. These findings infer the possibilities of antigenic drift, ensuing in compatibility of current vaccine for double mutants’ Indian strain. This double mutant strain seems to be a major burden for the available vaccine that could reduce the vaccine efficacy drastically and so may increase the chances of re-infection. However, more research is still needed to explicate the exact consequences of the double-mutant strain of SARS-CoV-2.

