## Supplementary File1 for "Bioinformatics analysis of SARS-CoV-2 RBD mutant variants and insights into antibody and ACE2 receptor binding"

Molecular interactions of antibody CR3022 with 7 different structurally changed RBD variants. Hydrogen bonds are depicted in green dashed lines and hydrophobic contacts are indicated by the red brick.

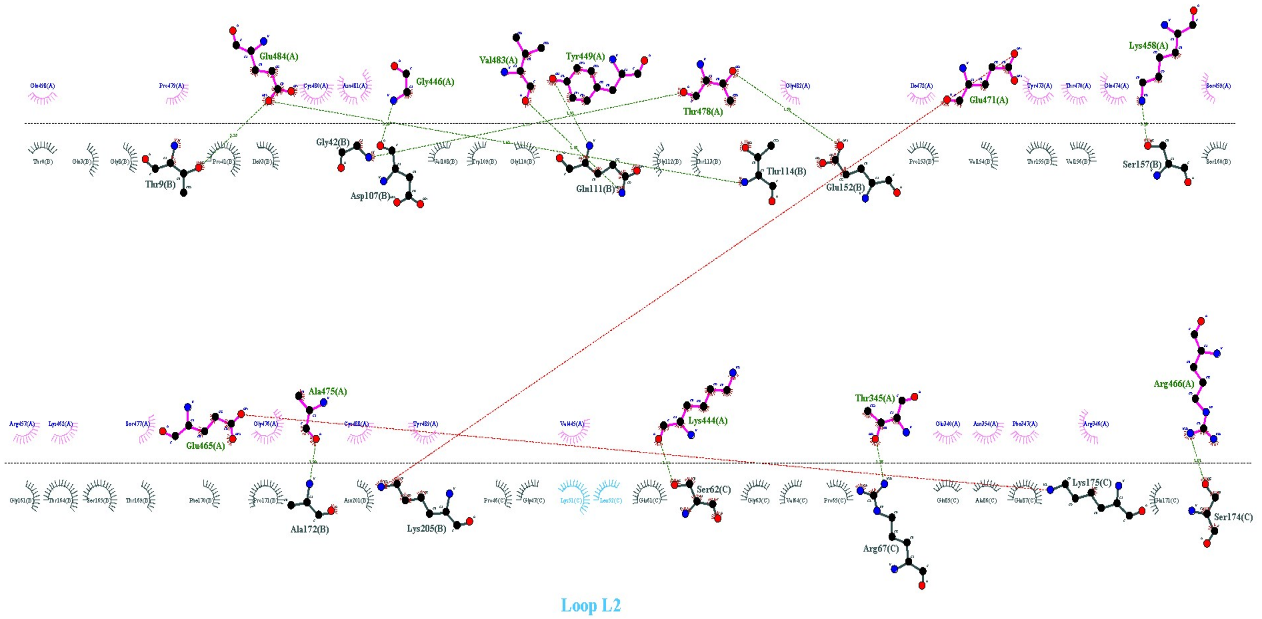


Spike wild

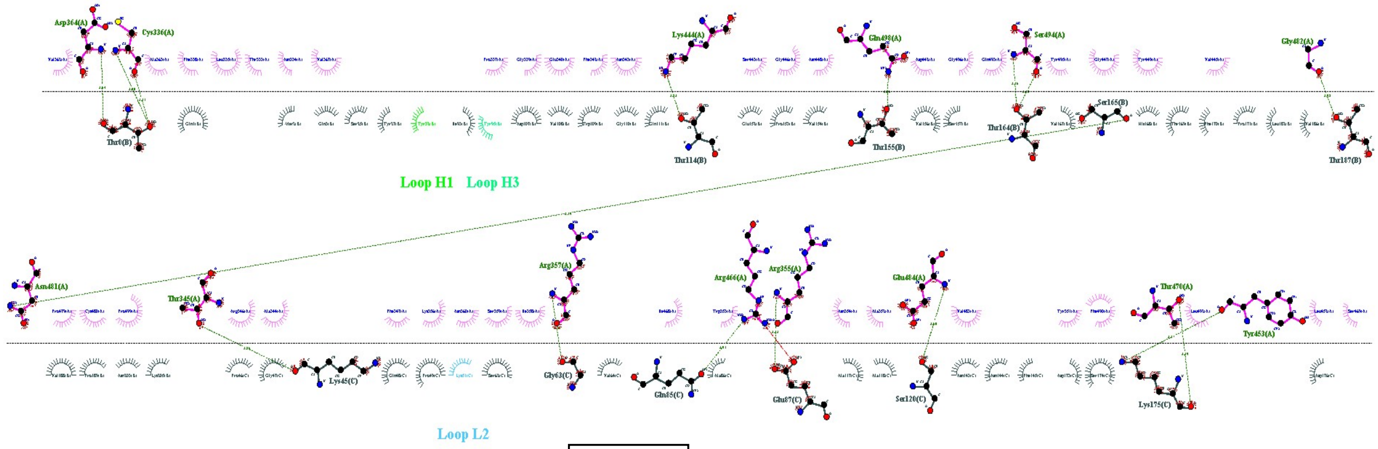


F486L

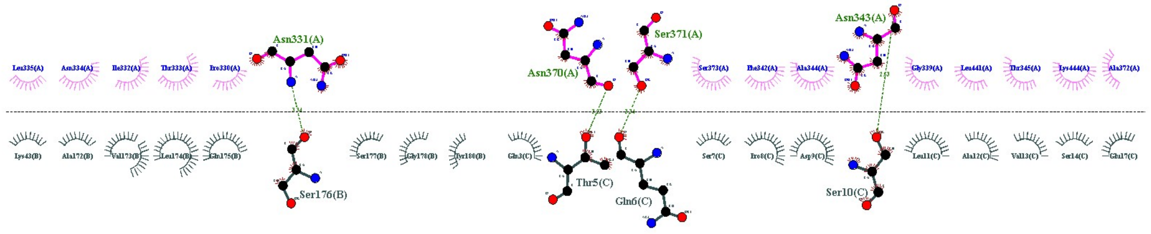


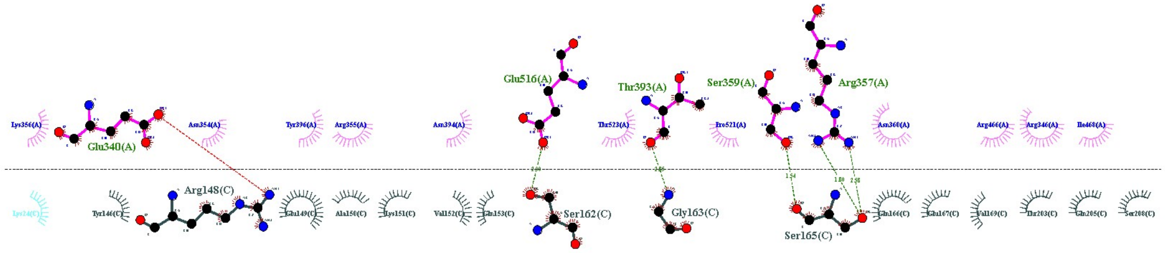


Loop L1

Q493N

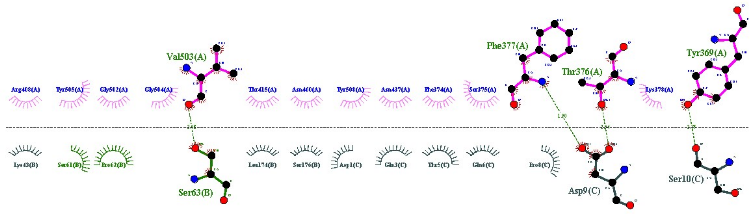


Loop H2

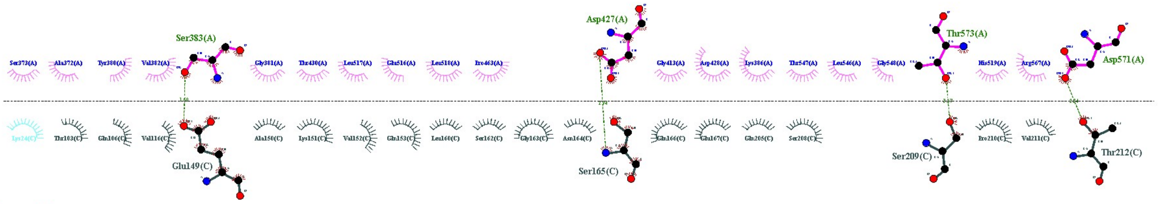


Loop L1

Double mutant

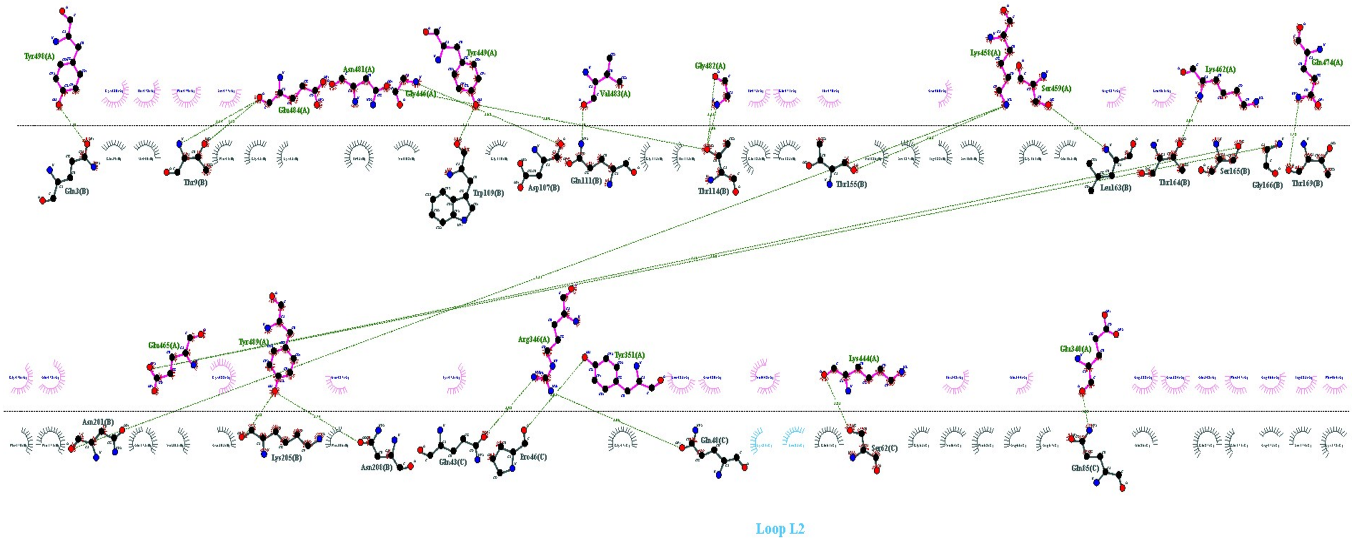
Loop L2

Q498Y

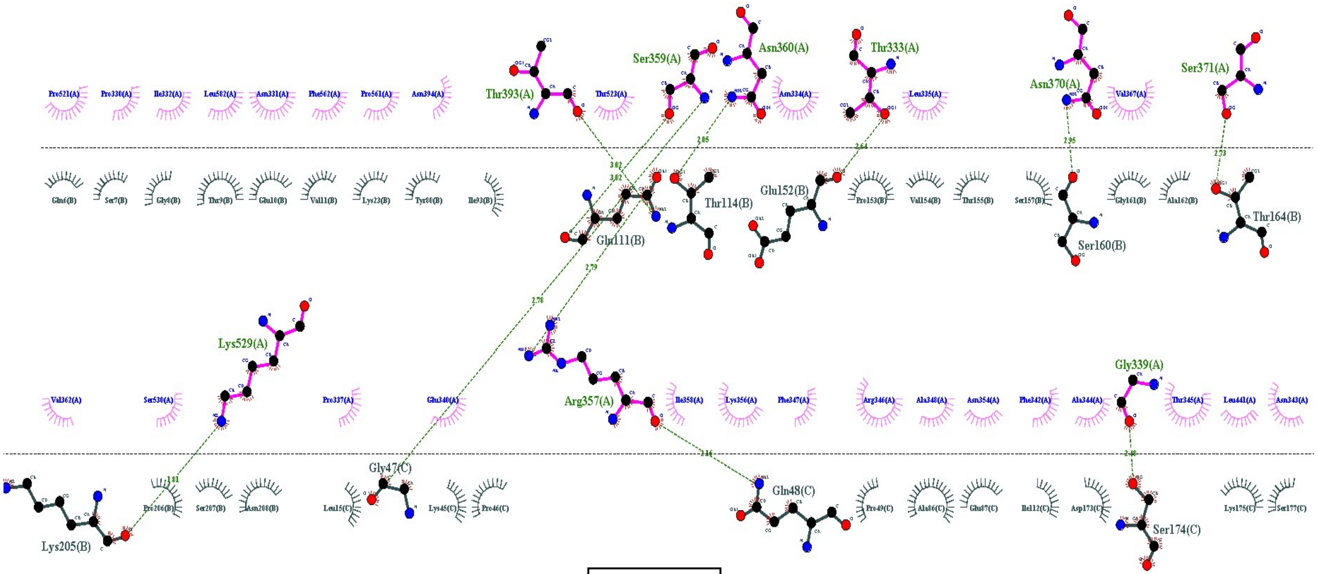
Ser359(A)
Asn360(A)
Thr333(A)
Asn370(A)
Ser371(A)
Thr393(A)
Thr114(B)
Glu152(B)
Glu111(B)
Ser160(B)
Thr164(B)
Lys529(A)
Gly339(A)
Arg357(A)
Gly47(C)
Gln48(C)
Ser174(C)
Lys205(B)

R408I

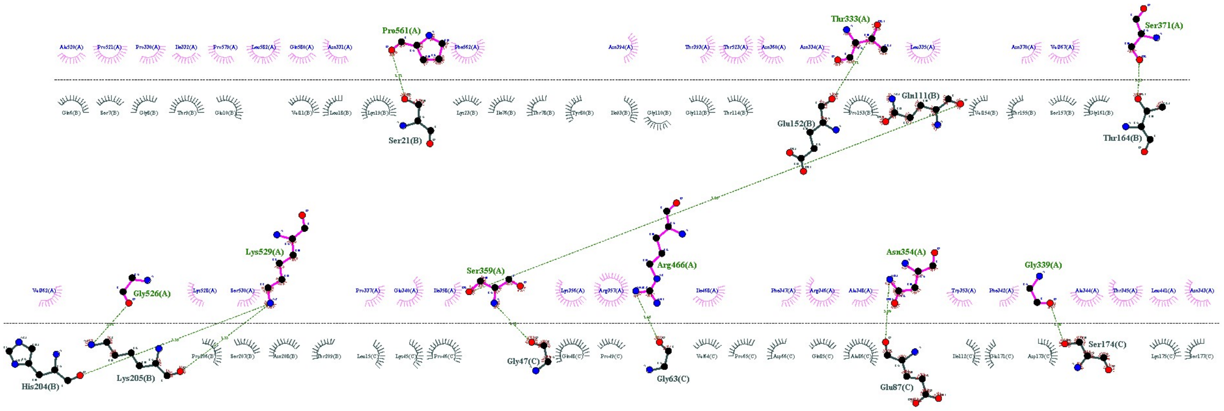
Pro561(A)
Thr333(A)
Ser371(A)
Gln111(B)
Glu152(B)
Ser21(B)
Thr164(B)
Lys529(A)
Gly526(A)
Ser359(A)
Arg466(A)
Asn354(A)
Gly339(A)
His204(B)
Lys205(B)
Gly47(C)
Gly63(C)
Glu87(C)
Ser174(C)

L455Y

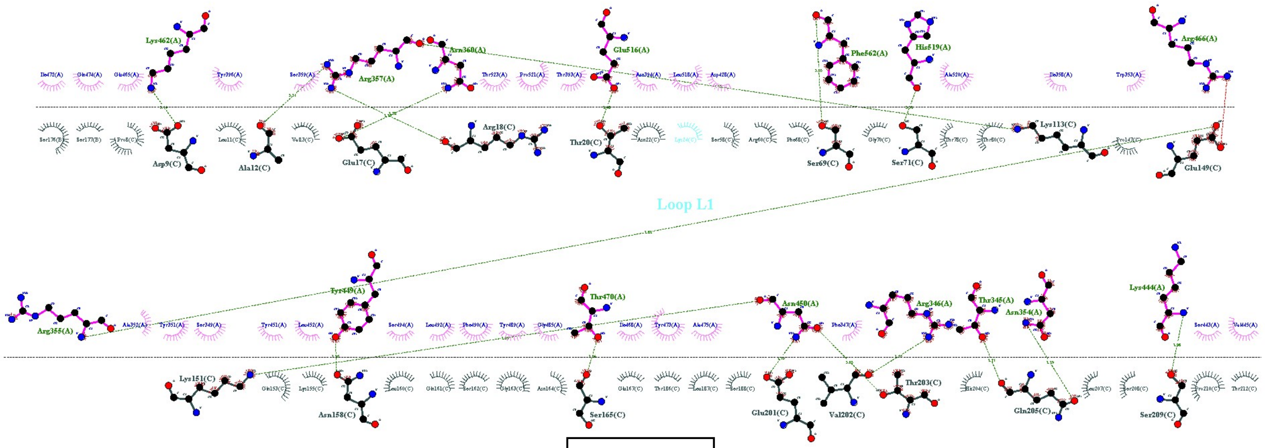
Lys462(A)
Arg357(A)
Asn360(A)
Glu516(A)
Phe562(A)
His519(A)
Arg466(A)
Asp9(C)
Ala12(C)
Glu17(C)
Arg18(C)
Thr20(C)
Ser69(C)
Ser71(C)
Lys113(C)
Glu149(C)
Loop L1
Arg355(A)
Tyr449(A)
Thr470(A)
Asn450(A)
Arg346(A)
Thr345(A)
Asn354(A)
Lys444(A)
Lys151(C)
Asn158(C)
Ser165(C)
Glu201(C)
Val202(C)
Thr203(C)
Gln205(C)
Ser209(C)

K417G

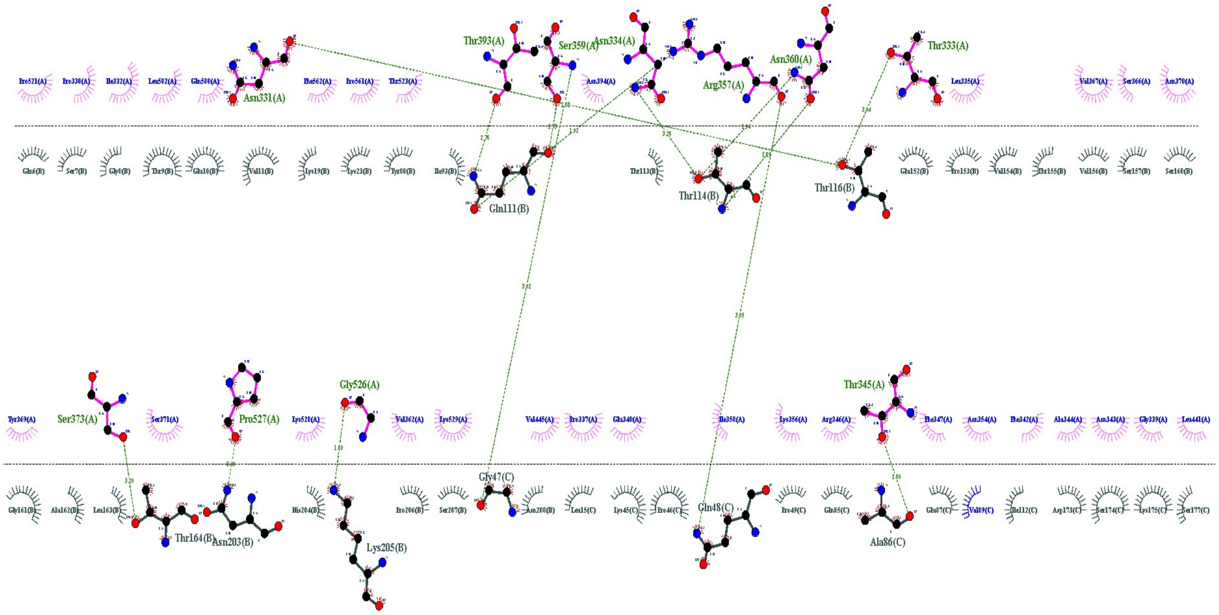


**Loop L3**

E484K

### ACE2 – RBD Variants

Molecular interaction of ACE2 (6ACG) with 7 different RBD Variants which have structural changes in RBD region. Hydrogen bonds showed in green dashed lines and hydrophobic contacts indicated by the red brick

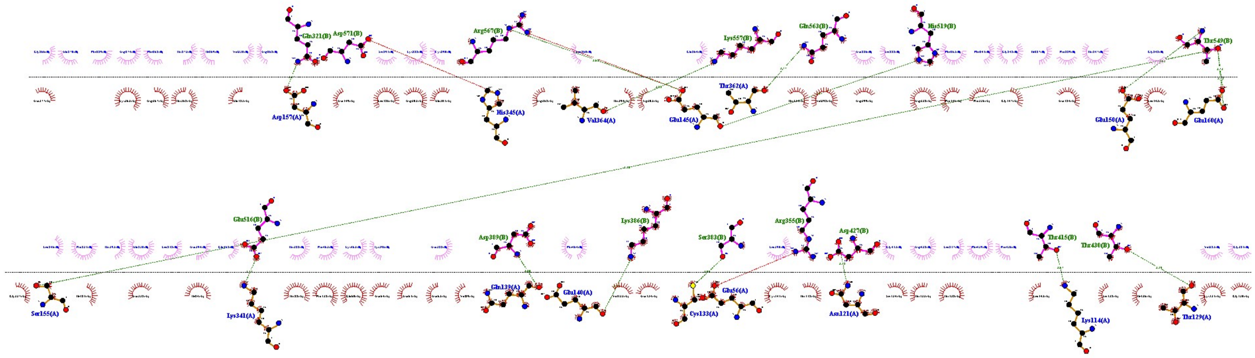

Spike Wild

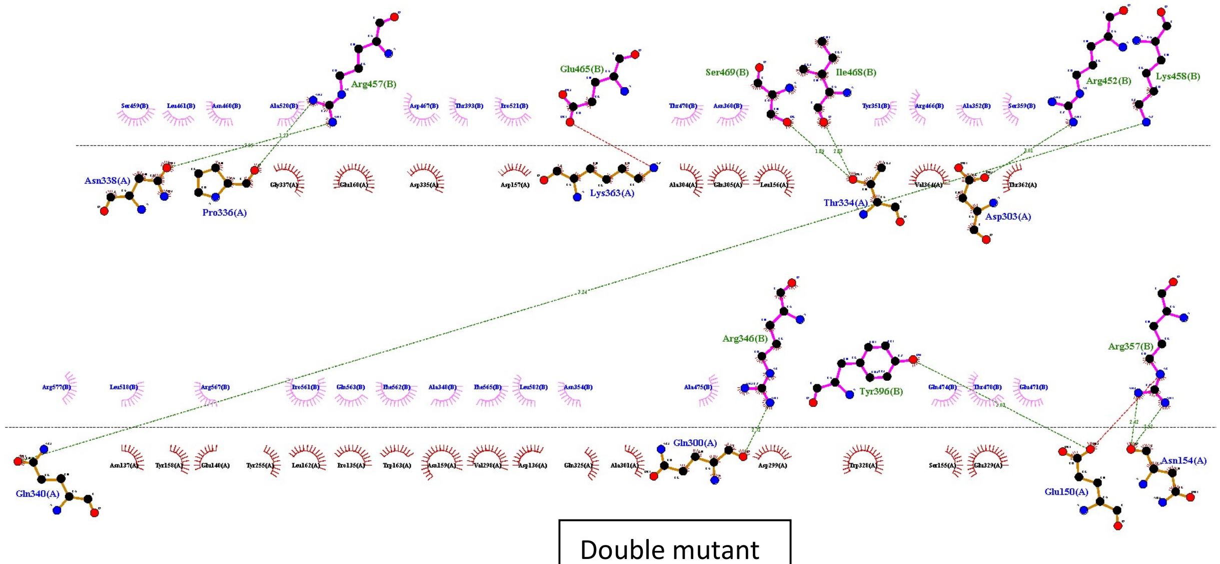


Double mutant

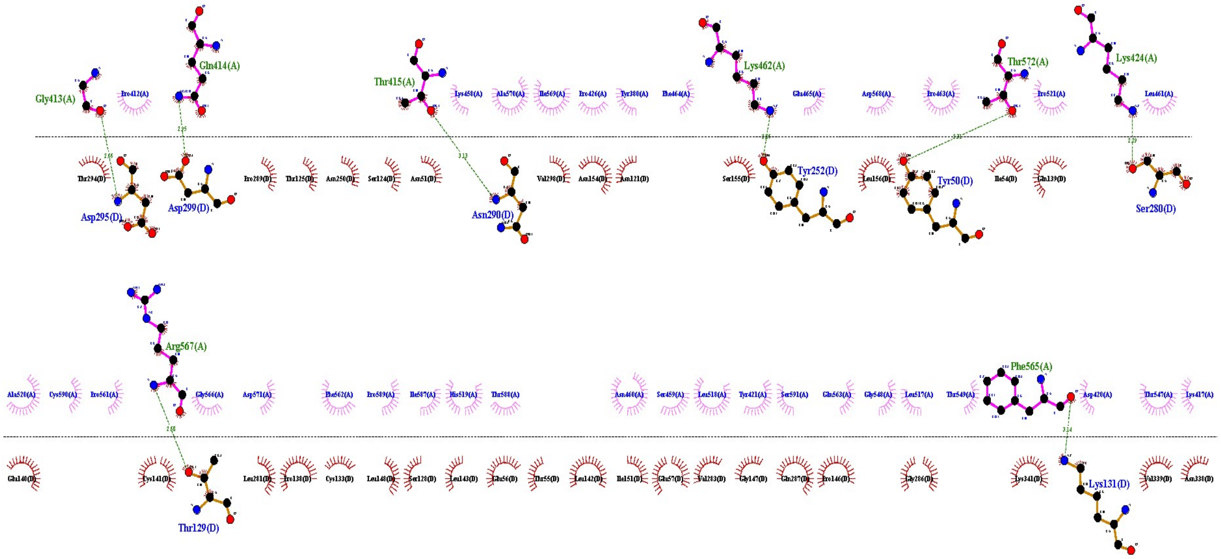
Gly413(A)
Gln414(A)
Thr415(A)
Lys462(A)
Thr572(A)
Lys424(A)
Asp295(D)
Asp299(D)
Asn290(D)
Tyr252(D)
Tyr50(D)
Ser280(D)
Arg567(A)
Phe565(A)
Thr129(D)
Lys131(D)

E484k

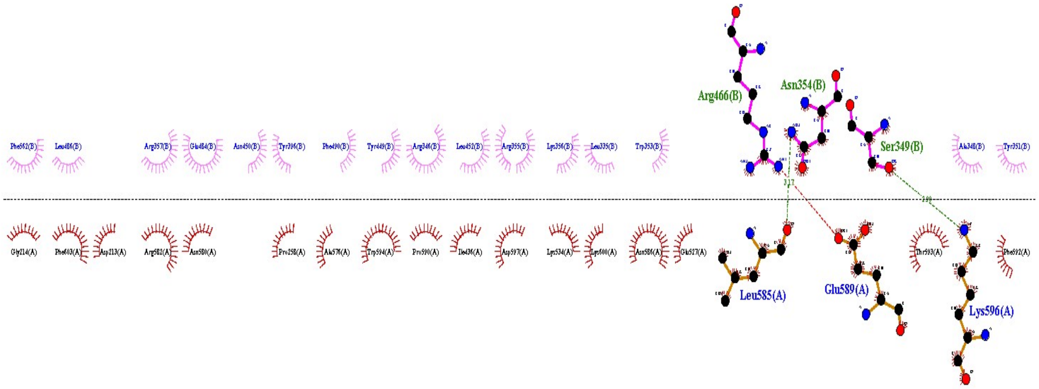
Arg466(B)
Asn354(B)
Ser349(B)
Leu585(A)
Glu589(A)
Lys596(A)

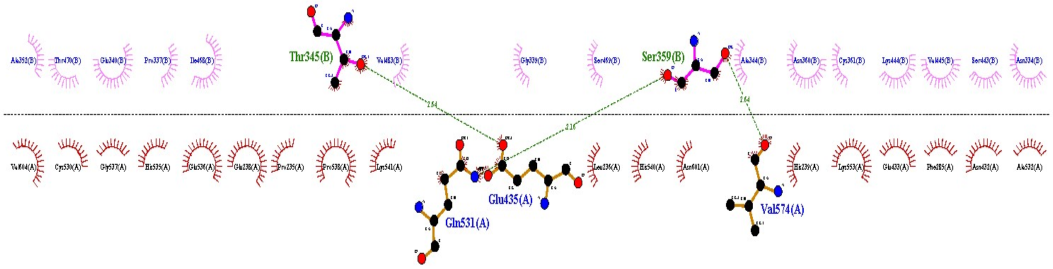
Thr345(B)
Ser359(B)
Glu435(A)
Gln531(A)
Val574(A)

F486L

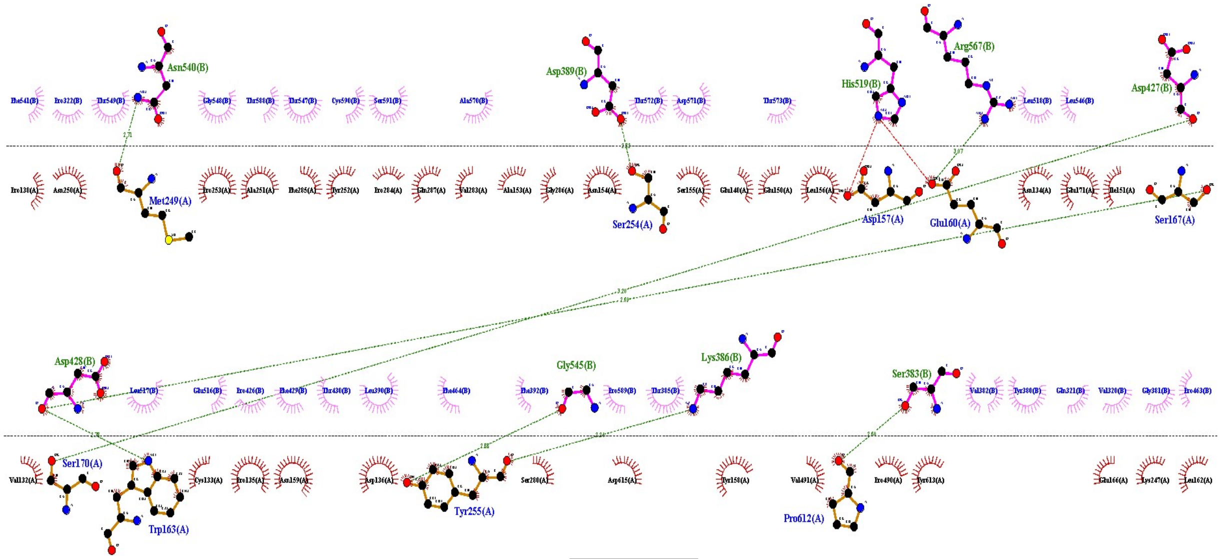

K417G

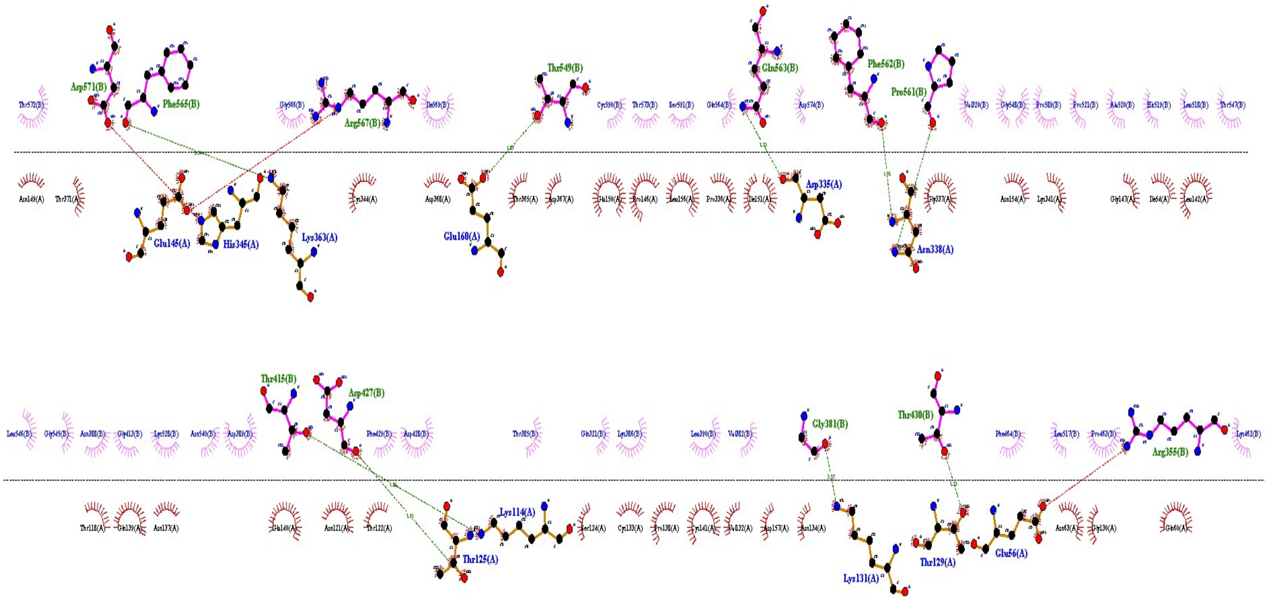

L455Y

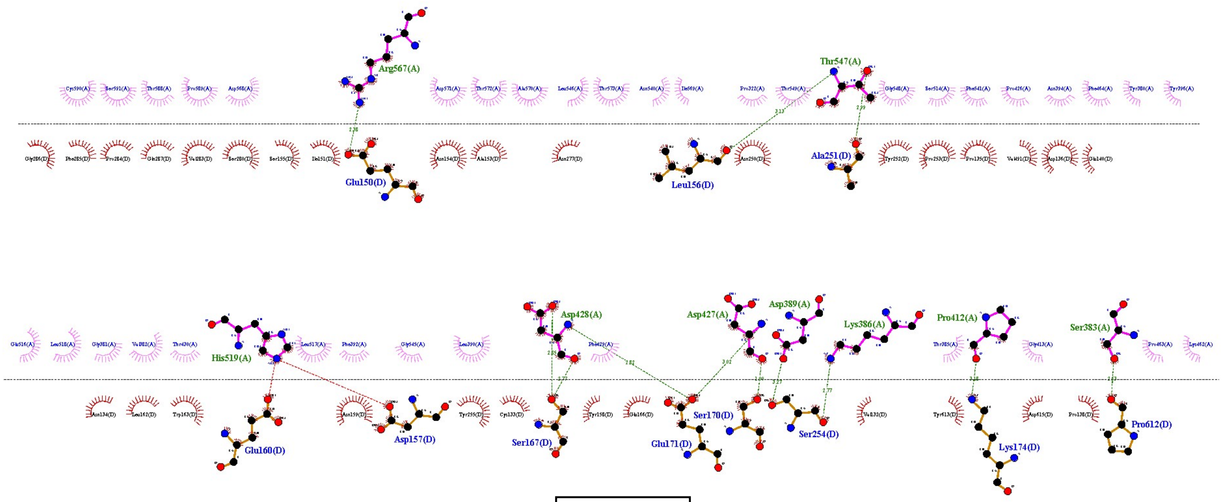
Arg567(A)
Thr547(A)
Glu150(D)
Leu156(D)
Ala251(D)
Asp428(A)
Asp427(A)
Asp389(A)
Lys386(A)
Pro412(A)
Ser383(A)
His519(A)
Glu160(D)
Asp157(D)
Ser167(D)
Ser170(D)
Glu171(D)
Ser254(D)
Lys174(D)
Pro612(D)

Q493N

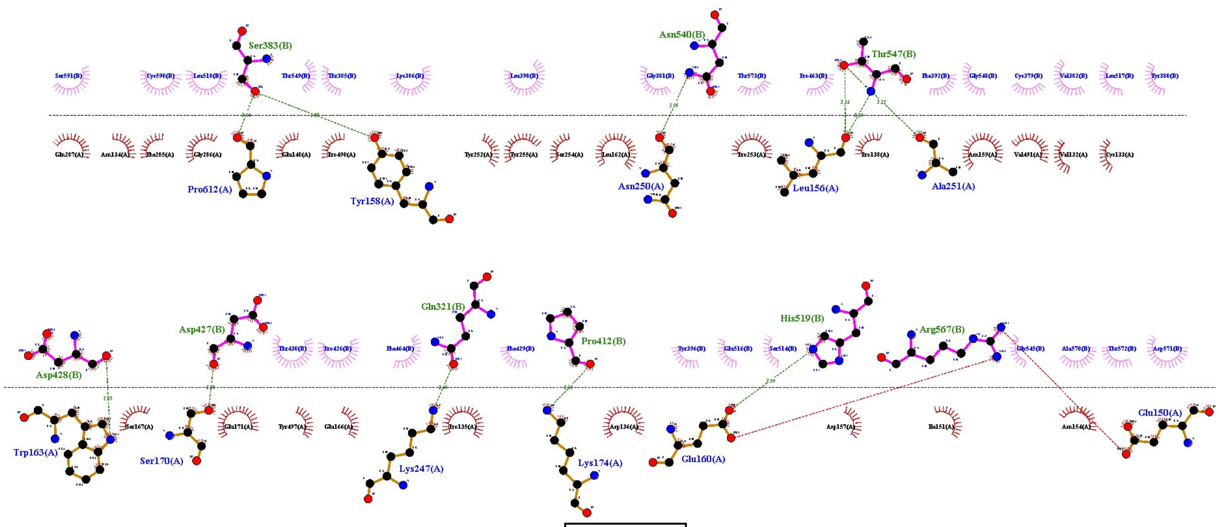
Ser383(B)
Asn540(B)
Thr547(B)
Pro612(A)
Tyr158(A)
Asn250(A)
Leu156(A)
Ala251(A)
Gln321(B)
Asp427(B)
Pro412(B)
His519(B)
Arg567(B)
Asp428(B)
Trp163(A)
Ser170(A)
Lys247(A)
Lys174(A)
Glu160(A)
Glu150(A)

R408I
